# Discovery of SARS-CoV-2 papain-like protease inhibitors through a combination of high-throughput screening and FlipGFP-based reporter assay

**DOI:** 10.1101/2021.03.15.435551

**Authors:** Zilei Xia, Michael Dominic Sacco, Chunlong Ma, Julia Alma Townsend, Naoya Kitamura, Yanmei Hu, Mandy Ba, Tommy Szeto, Xiujun Zhang, Xiangzhi Meng, Fushun Zhang, Yan Xiang, Michael Thomas Marty, Yu Chen, Jun Wang

## Abstract

The papain-like protease (PL^pro^) of SARS-CoV-2 is a validated antiviral drug target. PL^pro^ is involved in the cleavage of viral polyproteins and antagonizing host innate immune response through its deubiquitinating and deISG15ylating activities, rendering it a high profile antiviral drug target. Through a FRET-based high-throughput screening, several hits were identified as PL^pro^ inhibitors with IC_50_ values at the single-digit micromolar range. Subsequent lead optimization led to potent inhibitors with IC_50_ values ranging from 0.56 to 0.90 µM. To help prioritize lead compounds for the cellular antiviral assay against SARS-CoV-2, we developed the cell-based FlipGFP assay that is suitable for quantifying the intracellular enzymatic inhibition potency of PL^pro^ inhibitors in the BSL-2 setting. Two compounds selected from the FlipGFP-PL^pro^ assay, Jun9-53-2 and Jun9-72-2, inhibited SARS-CoV-2 replication in Caco-2 hACE2 cells with EC_50_ values of 8.89 and 8.32 µM, respectively, which were 3-fold more potent than GRL0617 (EC_50_ = 25.1 µM). The X-ray crystal structures of PL^pro^ in complex with GRL0617 showed that binding of GRL0617 to SARS-CoV-2 induced a conformational change in the BL2 loop to the more closed conformation. Overall, the PL^pro^ inhibitors identified in this study represent promising starting points for further development as SARS-CoV-2 antivirals, and FlipGFP-PL^pro^ assay might be a suitable surrogate for screening PL^pro^ inhibitors in the BSL-2 setting.

## INTRODUCTION

The COVID-19 pandemic has led to 120,105,958 confirmed cases and 2,657,629 deaths as of March 15, 2021, rendering it the worst pandemic since the 1918 Spanish flu. The etiological agent of COVID-19 is SARS-CoV-2, a single-stranded positive sense RNA virus that belongs to the beta coronavirus family. Two additional coronaviruses within the same family, the SARS-CoV, and MERS-CoV, have caused epidemics in humans with mortality rates of 9.6% and 34.3%, respectively. Although SARS-CoV-2 has a lower mortality rate of 2.2% compared to SARS-CoV and MERS-CoV, it has led to far greater death tolls due to its higher transmission. SARS-CoV-2 differs from SARS-CoV and MERS-CoV in that it has a long incubation time after the initial infection (∼ 2 weeks), and a large percentage of infected patients continue to shed the virus while being asymptomatic, presenting a daunting task for surveillance and containment.

Three mRNA vaccines developed by Pfizer/BioNtech, Moderna, and Johnson and Johnson have been approved by FDA in the United States. For small molecule antivirals, remdesivir received FDA approval on October 22, 2020. Although the polymerase of SARS-CoV-2 has proof-reading function, it continues to mutate at a rate about 10^−6^ per site per cycle.^1^ Several mutant viruses have already emerged and widely circulated among humans since the beginning of the pandemic.^2^ Therefore, there is a dire need of additional antivirals with a novel mechanism of action. Antivirals are not substituents of vaccines, but rather an important complement to help combat the pandemic. Among the viral proteins that have been actively pursued as SARS-CoV-2 antiviral drug targets, the main protease (M^pro^) and papain-like protease (PL^pro^) are the most promising ones.^3,4^ Both M^pro^ and PL^pro^ are involved in the proteolytic digestion of the viral polyproteins pp1a and pp1ab, yielding individual functional viral proteins for the replication complex formation. PL^pro^ cleaves at three sites with the recognition sequence “LXGG XX”.^5^ PL^pro^ has been shown to play additional roles in dysregulating host immune response and impairing the host type I interferon antiviral effect through its deubiquitinating and deISG15ylating (interferon-induced gene 15) activities, respectively.^6-8^ SARS-CoV-2 PL^pro^ cleaves ISG15 and polyubiquitin modifications from cellular proteins, and inhibition of PL^pro^ led to the accumulation of ISG15-conjugates and poly-ubiquitin-conjugates.^9^ While SARS-CoV PL^pro^ prefers ubiquitinated substrates, the SARS-CoV-2 PL^pro^ prefers the ISGlyated proteins as substrates.^6-8^ PL^pro^ is part of a membrane anchored multi-domain protein named non-structural protein 3 (nsp-3), an essential component of the replicase-transcriptase complex. The multi-functional roles of SARS-CoV-2 PL^pro^ render it a prominent antiviral drug target. A substantial morbidity and mortality associated with COVID-19 infection is caused by cytokine storm,^10^ and suppressing host immune response using dexamethasone and baricitinib has been shown to provide therapeutic benefits in the treatment of severe infections.

A significant progress has been made in developing SARS-CoV-2 M^pro^ inhibitors^3,4,11-13^ and the Pfizer compound PF-07304814 currently in phase 1 clinical trial.^14^ In comparison, PL^pro^ represents a more challenging drug target, and GRL0617 remains one of the most potent PL^pro^ inhibitors reported to date despite several high-throughput screening and medicinal chemistry optimization campaigns.^8,15^ GRL0617 was originally developed against SARS-CoV PL^pro^.^16^ As SARS-CoV-2 and SARS-CoV PL^pro^ share a sequence identity of 83% and similarity of 90%, GRL0617 was also repurposed for SARS-CoV-2 PL^pro^ and it was reported to inhibit SARS-CoV-2 PL^pro^ with IC_50_ values of around 2 µM and the SARS-CoV-2 viral replication with EC_50_ values around 20 µM from multiple studies.^8,9,15,17^

In this study, we report our progress in developing novel SARS-CoV-2 PL^pro^ inhibitors. Using the FRET-based enzymatic assay, we conducted a high-throughput screening against the Enamine 50K diversity compound library and identified two hits Jun9-13-7 and Jun9-13-9 with single-digit micromolar IC_50_ values. Upon validating the hits using differential scanning fluorimetry (DSF) and native mass spectrometry binding assays, a focused library of structural analogs were designed and tested to dissect the structure-activity relationship (SAR) studies. Several compounds have been identified to inhibit SARS-CoV-2 PL^pro^ with submicromolar potency. To prioritize lead compounds for the antiviral assay against SARS-CoV-2, we developed the FlipGFP assay that is suitable for quantifying the intracellular activity of PL^pro^ inhibitors in the BSL2 setting. Two compounds selected from the FlipGFP PL^pro^ assay, Jun9-53-2 and Jun9-72-2, showed 3-fold improvement than GRL0617 in the cellular antiviral assay against SARS-CoV-2. X-ray crystal structures showed that binding of GRL0617 to the wild-type (WT) SARS-CoV-2 PL^pro^ induced a conformational change in the BL2 loop to the more closed conformation. Overall, the SARS-CoV-2 PL^pro^ reported herein represent promising hits for further development as SARS-CoV-2 antivirals, and FlipGFP PL^pro^ assay might be useful in testing the cellular activity of PL^pro^ inhibitors in the BSL-2 setting.

## RESULTS AND DISCUSSION

### Expression and characterization of SARS-CoV-2 PL^pro^

Two constructs of SARS-CoV-2 PL^pro^ were expressed in E. Coli, one with Hig-tag (PL^pro^-His) and another without the tag (PL^pro^). To profile the proteolytic activity of PL^pro^ in cleaving the viral polyprotein, we developed a FRET based enzymatic assay with the peptide substrate 4-((4-(dimethylamino)phenyl)azo)benzoic acid (Dabcyl)– FTLRGG/APTKV–5-[(2-aminoethyl)amino]naphthalene-1-sulfonic acid (Edans), which corresponds to the nsp2-nsp3 junction from the SARS-CoV-2 polyprotein. The enzymatic activity k_cat_/K_m_ of PL^pro^-His and PL^pro^ were 340 M^-1^S^-1^ and 255 M^-1^S^-1^ (Table S1), respectively, which were similar to previous reports,^8,17^ suggesting both constructs were enzymatically active. The SARS-CoV-2 PL^pro^ was also reported to have deubiquitinating and deISGylating activities. Accordingly, we characterized the deubiquitinating and deISGylating activities of SARS-CoV-2 PL^pro^ using the Ub-AMC and ISG-AMC substrates, respectively, in the enzymatic assay. It was found that the SARS-CoV-2 PL^pro^ is more efficient in cleaving the ubiquitin (Ub) and ISG15 (ISG) modifications than the viral polyprotein, with k_cat_/K_m_ values of 1,070 and 1.67×10^5^ M^-1^S^-1^ (Table S2), respectively. This substrate preference is in agreement with results reported by Gal et al,^17^ and SARS-CoV PL^pro^ was also reported to have similar substrate preference.^18^ Significantly, the deISGylating activity is 156-fold higher than the deubiquitinating activity, which is consistent with previous reports that the SARS-CoV-2 PL^pro^ prefers ISG15 over ubiquitin as a substrate.^5-9^

### High-throughput screening of the Enamine 50K diversity library against the SARS-CoV-2 PL^pro^ and hit validation

The HTS assay was optimized in 384-well plates using the FRET substrate, which gave a Z’ factor of 0.668 with a signal to noise ratio (S/B) of 11.2, indicating that this was a robust assay (Fig. 1). We then performed the HTS against the enamine library, which consists of 50,240 structurally diverse compounds. GRL0617 was included as a positive control.

**Fig 1.**
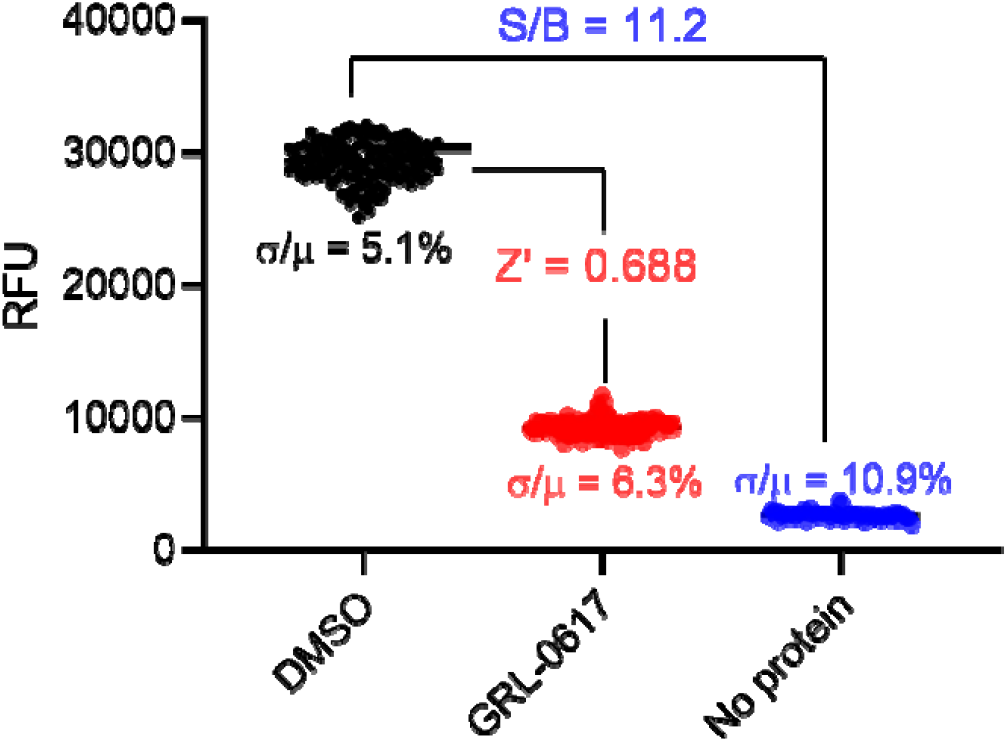
384-well high-throughput screening assay with SARS CoV-2 PL^pro^. In this representative control plate (column 1 to 20 with PL^pro^ protein, column 21 to 24 with buffer only; 1 µl DMSO was added to column 1 to 10 and 21 to 24; 1 µl GRL0617 was added to column 11 to 20), the signal to base ratio (S/B) is 11.2, and the calculated Z’ factor is 0.688.

Hits showing more than 50% inhibition were repurchased from Enamine and titrated in the FRET-based enzymatic assay to determine the IC_50_ values (Fig. 2A and Table S3). In parallel, differential scanning fluorimetry (DSF) assay was performed as a secondary assay to characterize the binding of the hits with SARS-CoV-2 PL^pro^ (Fig. 2B and Table S3). The most potent two hits Jun9-13-7 and Jun9-13-9 (Fig. 2C) had IC_50_ values of 7.29 ± 1.03 and 6.67 ± 0.05 µM, respectively. Compounds Jun9-13-7 and Jun9-13-9 also increased the thermal stability of SARS-CoV-2 PL^pro^ by 2.98 ± 0.09 and 2.18 ± 0.29 °C (Table S3), which is consistent with their enzymatic inhibition. In comparison, GRL0617 had an IC_50_ value of 2.05 ± 0.12 µM, and increased the protein stability by 3.52 ± 0.27 °C in the DSF assay (Table S3). The potency of GRL0617 in inhibiting SARS-CoV-2 PL^pro^ from our study is consistent with recent reports.^6-9^ The rest of the hits had weak enzymatic inhibition (IC_50_ > 10 µM) and showed marginal binding to PL^pro^, therefore they were not further pursued (Table S3). Both compound Jun9-13-7 and Jun9-13-9 also inhibit the deubiquitinating and deISGylating activities with IC_50_ values ranging from 4.93 to 12.5 µM (Fig. 2D and Table S4). In contrast, neither of these two compounds inhibited the SARS-CoV-2 M^pro^ at up to 200 µM (fig. S2), suggesting the inhibition of SARS-CoV-2 PL^pro^ by these two compounds are specific. The binding of Jun9-13-7 and Jun9-13-9 to SARS-CoV-2 PL^pro^ was further characterized using the native mass spectrometry (Fig. 2E). It was shown that both Jun9-13-7 and Jun9-13-9 showed dose-dependent binding to PL^pro^ with binding stoichiometry of one drug per PL^pro^, similar to the positive control GRL0617. Enzymatic kinetic studies showed that compounds Jun9-13-7 and Jun9-13-9 are non-covalent inhibitors with Ki values of 3.96 and 2.10 µM, respectively (fig. S3). The Lineweaver-Burk plots yielded an intercept at the Y-axis, suggesting that both compounds are competitive inhibitors similar to GRL0617 (fig. S3).

**Fig 2.**
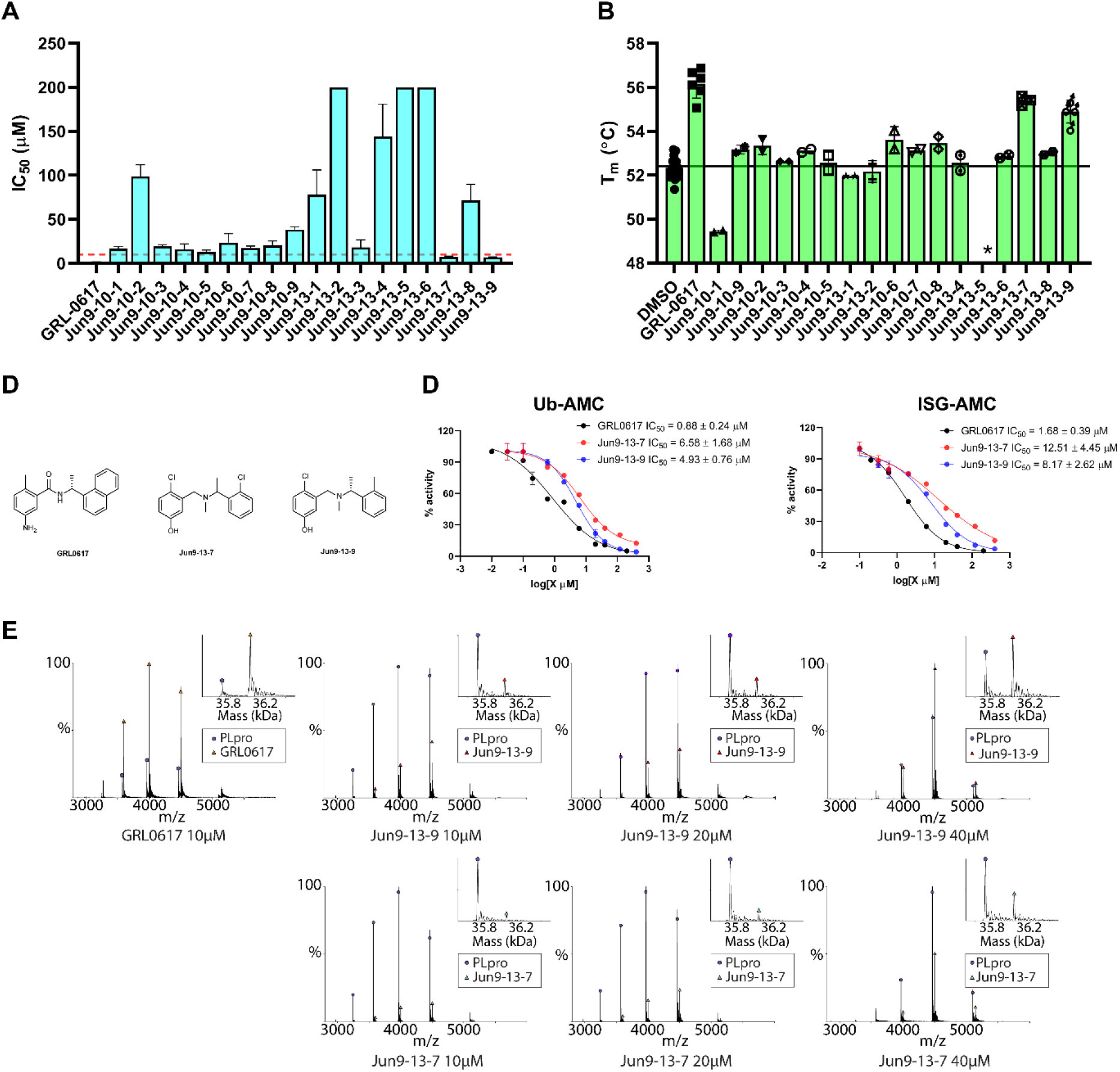
HTS and hit validation of SARS-CoV-2 PL^pro^ inhibitors. (**A**) IC_50_ values of the screening hits in the FRET-based enzymatic assay, the red line indicates the IC_50_ = 10 µM. (**B**) Differential scanning fluorimetry assay of the screening hits in stabilizing the SARS-CoV-2 PL^pro^. (**C**) Chemical structures of GRL0617, Jun9-13-9 and Jun9-13-7. (**D**) Inhibitory activity of Jun9-13-7 and Jun9-13-9 against SARS-CoV-2 PL^pro^ using Ub-AMC and ISG-AMC substrates. (**E**) Native MS binding assay of Jun9-13-9 and Jun9-13-7 to SARS-CoV-2 PL^pro^.

### Lead optimization of SARS-CoV-2 PL^pro^ inhibitors

To further optimize the enzymatic inhibition of Jun9-13-7 and Jun9-13-9, 13 structural analogs were purchased from Enamine (Fig. 3A) and 34 compounds were synthesized (Fig. 3B) to elucidate the structure-activity relationships (SAR). It was found that the hydroxyl substation at the left phenyl ring is critical for the activity as methylation led to significant loss of enzymatic inhibition (Jun9-13-9 vs Jun9-25-4). The methylene linker and the methyl substation on the link are both important for the enzymatic inhibition (Jun9-13-9 vs Jun9-26-2). Similarly, the ortho-methyl or chloride substation on the right phenyl ring is critical for the activity (Jun9-13-7 vs Jun9-29-5; Jun9-13-7 vs Jun9-13-4). Next, guided by this initial SAR results, 34 analogs were designed and synthesized (Fig. 3B). Nine compounds had IC_50_ values less than 1 µM including Jun9-75-4 (IC_50_ = 0.62 µM), Jun9-85-1 (IC_50_ = 0.66 µM), Jun9-84-3 (IC_50_ = 0.67 µM), Jun9-87-1 (IC_50_ = 0.87 µM), Jun9-72-2 (IC_50_ = 0.67 µM), Jun9-87-2 (IC_50_ = 0.90 µM), Jun9-87-3 (IC_50_ = 0.80 µM), Jun9-75-5 (IC_50_ = 0.56 µM), and Jun9-53-2 (IC_50_= 0.89 µM). Among them, Jun9-75-4 was the most potent PL^pro^ inhibitor with an IC_50_ of 0.62 µM, a 10-fold increase compared to Jun9-13-9 (IC_50_ = 6.67 µM). Jun9-75-4 is also 3-fold more potent than GRL0617 (IC_50_ = 2.05 ± 0.12 µM), representing one of the most potent PL^pro^ inhibitors reported to date.

**Fig 3.**
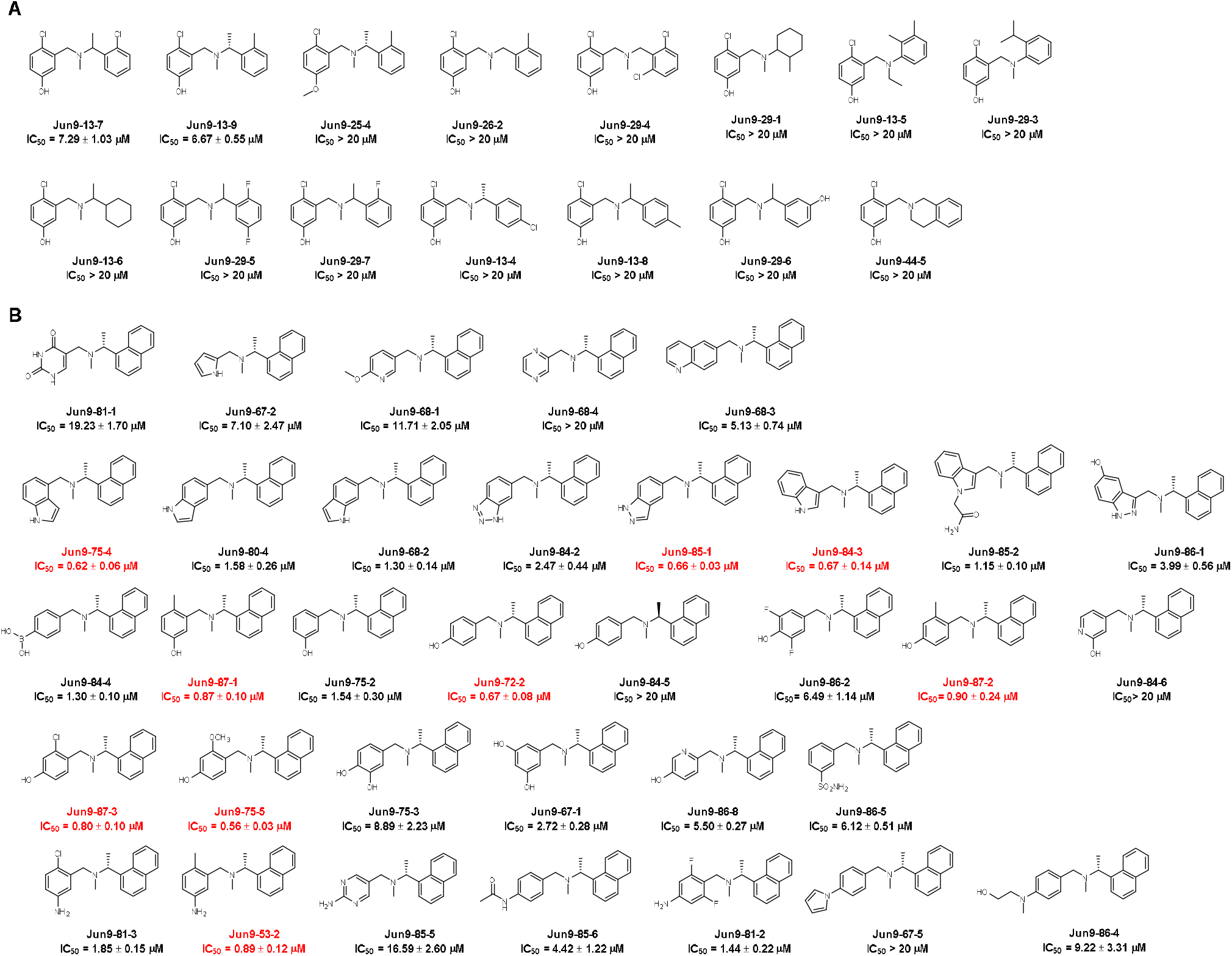
SAR of SARS-CoV-2 PL^pro^ inhibitors. (**A**) Analogs purchased from Enamine. (**B**) Synthetic compounds designed based on the SAR results. Potent compounds with IC_50_ values less than 1 µM were highlighted in red.

### Development of FlipGFP assay for testing the cellular activity of SARS-CoV-2 PL^pro^ inhibitors

One of the challenges in SARS-CoV-2 antiviral drug discovery is that SARS-CoV-2 is a biosafety level 3 (BSL3) pathogen, which limits the number of drug candidates that can be screened. To help prioritize lead compounds for the antiviral assay with infectious SARS-CoV-2, we developed a cell-based FlipGFP assay for SARS-CoV-2 PL^pro^ that is suitable for testing the intracellular activity of PL^pro^ inhibitors in the BSL2 setting. The advantages of cell-based PL^pro^ assay over the FRET-based enzymatic assay include but not limited to: 1) it can eliminate compounds that either cytotoxic or membrane impermeable; 2) In contrast to the standard FRET-based assay, PL^pro^ cleaves the substrate in the cell cytoplasm in the FlipGFP assay, which is a close mimetic of the viral polyprotein cleavage by PL^pro^ in virus-infected cell. It is known that cysteine proteases are susceptible to re-dox active compounds as well as non-specific alkylating chemicals such as ebselen.^19,20^ The FlipGFP PL^pro^ assay is expected to rule out such promiscuous compounds as the substrate is cleaved under the reducing intracellular environment.

In the assay design, the 10th and 11th β-strands from the GFP protein were separated from the rest of the GFP β-barrel (β-strands 1-9) (Fig. 4A).^21,22^ The 10^th^ and 11^th^ β-strands were linked through the PL^pro^ cleavage site and a heterodimerized coiled coils E5/K5. In the absence of the PL^pro^, the 10^th^ and 11^th^ β-strands are conformationally restrained and unable to associate with the GFP β-barrel 1-9. When the cleavage site is digested by the PL^pro^, the 11^th^ β-strand can then flip its orientation and associate with GFP β-barrel 1-9 together with the 10^th^ β-strand, leading to restoration of the green fluorescence signal (Fig. 4A). A red fluorescent protein mCherry was included within the construct via a “self-cleaving” 2A peptide to act as the transfection control (Fig. 4B), and the normalized ratio of green fluorescence signal over red fluorescence signal is proportional to the enzymatic activity of PL^pro^. Cells transfected with FlipGFP-PL^pro^ but without the PL^pro^ showed no green fluorescence signal (Fig. 4C, 6^th^ row), suggesting host proteases are unable to cleavage the PL^pro^ substrate sequence, thereby eliminating the background signal interference. A strong green fluorescence signal was observed only when there was a match between the cleavage site and the corresponding protease (Fig. 4C, 2^nd^ and 7^th^ rows). Specifically, little or no GFP signal was observed when the cells were transfected with SARS-CoV-2 PL^pro^ and a construct containing either the TEV cleavage site (FlipGFP-TEV) (Fig. 4C, 4^th^ row) or the M^pro^ cleavage site (FlipGFP-M^pro^) (Fig. 4C, 3^rd^ row). Similarly, little or no GFP signal was observed when the cells were transfected with SARS-CoV-2 M^pro^ and a construct containing the PL^pro^ cleavage site (FlipGFP-PL^pro^) (Fig. 4C, 5^th^ row). In contrast, strong green fluorescence signals were observed when the cells were transfected with PL^pro^ and FlipGFP-PL^pro^ (Fig. 4C, 7^th^ row) or M^pro^ and FlipGFP-M^pro^ (Fig. 4C, 2^nd^ row).

**Fig 4.**
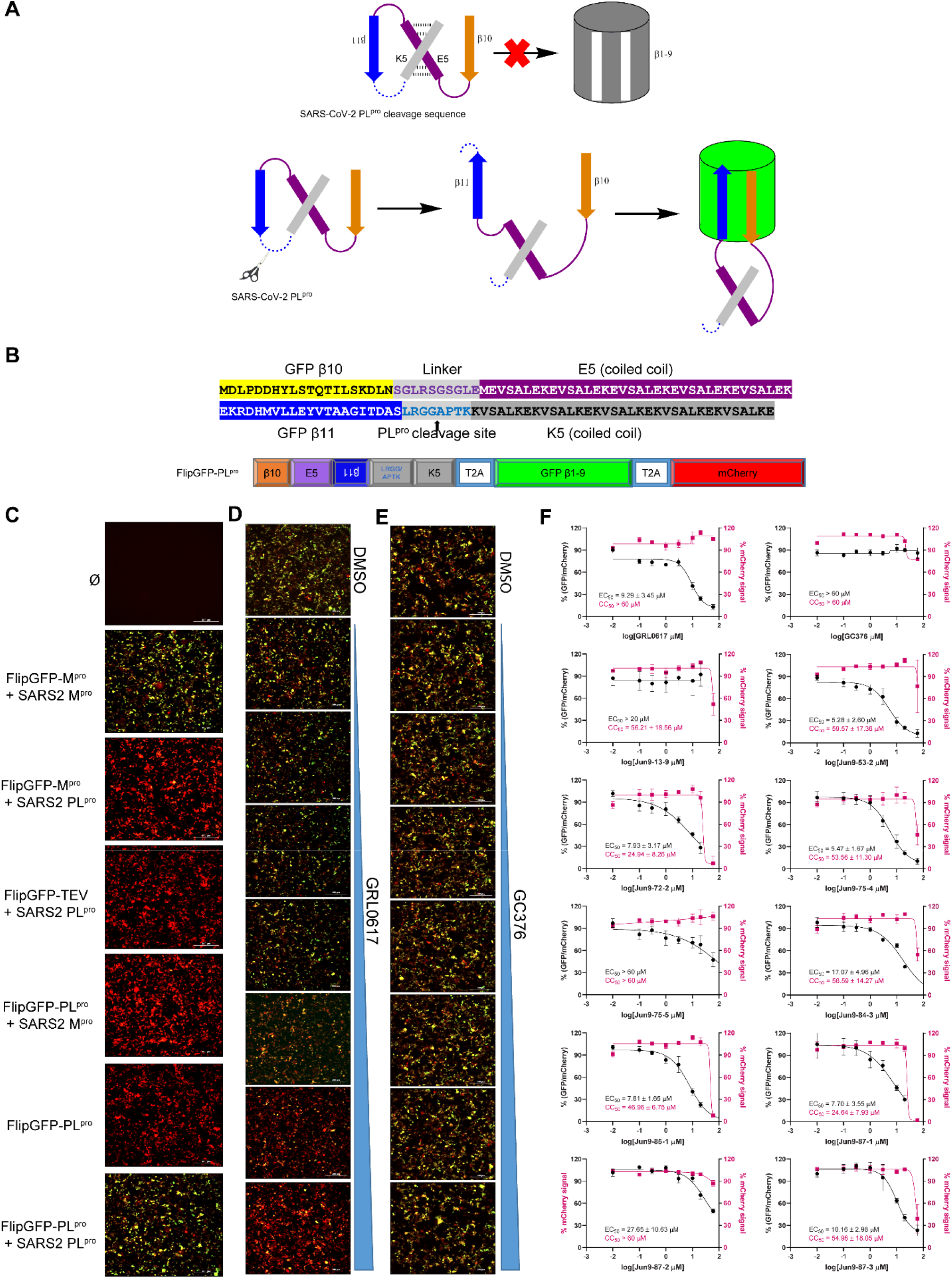
Development of cell-based FlipGFP assay for the quantification of the cellular activity of SARS-CoV-2 PL^Pro^ inhibitors. (**A**) Design principle for the cell- based FlipGFP assay. (**B**) Sequence of the flipped GFP β10-11 and cleavage site. (**C**) FlipGFP-PL^Pro^ assay development. 293T cells that were not transfected (∅), or transfected with FlipGFP-M^pro^ and SARS CoV-2 M^pro^ plasmids, or FlipGFP-M^pro^ and SARS CoV-2 PL^pro^ plasmids, or FlipGFP-TEV and SARS CoV-2 PL^pro^ plasmids, or FlipGFP-PL^pro^ and SARS CoV-2 M^pro^ plasmids, or FlipGFP-PL^pro^ plasmid alone, or FlipGFP-PL^pro^ and SARS CoV-2 PL^pro^ plasmids (details were described in method section). Representative images of FlipGFP-PL^pro^ assay with the positive control GRL0617 (**D**) and the negative control GC376 (**E**). (**F**) Dose-response inhibition PL^pro^ in the FlipGFP assay by ten compounds selected from the FRET-based enzymatic assay.

With the established assay condition, we then screened nine most potent PL^pro^ inhibitors with IC_50_ values less than 1 µM from the FRET-based enzymatic assay (Fig. 3). GRL0617 and GC376 were included as positive and negative controls, respectively. The initial hit Jun9-13-9 was also included. Compounds were added 3 h post transfection, and GFP and mCherry fluorescence signals were measured at 48 h post transfection using the Cytation 5 imager reader. A dose dependent decrease of the GFP signal was observed with increasing concentrations of GRL0617 (Fig. 4D), and quantification of the normalized GFP/mCherry ratio gave an EC_50_ value of 9.29 ± 3.45 µM. In contrast, GC376 had no effect on the intensity of green fluorescence signal (EC_50_ > 60 µM) (Fig. 4E), suggesting the FlipGFP assay is suitable for the screening of PL^pro^ inhibitors. Among the ten compounds tested in the cell-based FlipGFP assay, compounds Jun9-53-2, Jun9-72-2, Jun9-75-4, Jun9-85-1, and Jun9-87-1 had EC_50_ values less than 10 µM, while compounds Jun9-84-3, Jun9-87-2. And Jun9-87-3 were less active and had EC_50_ values between 10.16 to 17.07 µM. Compound Jun9-75-5 was not active (EC_50_ > 60 µM), despite its potent activity in the FRET-based enzymatic assay (IC_50_ = 0.56 µM).

### Cellular antiviral activity of PL^pro^ inhibitors against SARS-CoV-2

Based on the FlipGFP-PL^pro^ assay results, we selected two compounds Jun9-53-2 and Jun9-72-2 and tested their cellular antiviral activity against the infectious SARS-CoV-2 in two cell lines, the Caco-2 hACE2, and the Vero E6. The Caco-2 hACE2 cell line expresses endogenous TMPRSS2 and was transfected to overexpress hACE2, rending it a physiological relevant cell line model that is close to the human airway epithelial cells. GRL0617 was included as a positive control. It was found that all three compounds had more potent inhibition in Caco-2 hACE2 cells than the Vero E6 cells (Fig. 5). Consistent with the enzymatic inhibition results, Jun9-53-2 and Jun9-72-2 had more potent cellular antiviral activity than GRL0617 in the two cell lines tested. The EC_50_ values of GRL0617, Jun9-53-2, and Jun9-44-3, in inhibiting SARS-CoV-2 in Caco-2 hACE2 cell were 25.14 ± 7.58, 8.89 ± 3.28, and 8.32 ± 1.48 µM, respectively.

**Fig 5.**
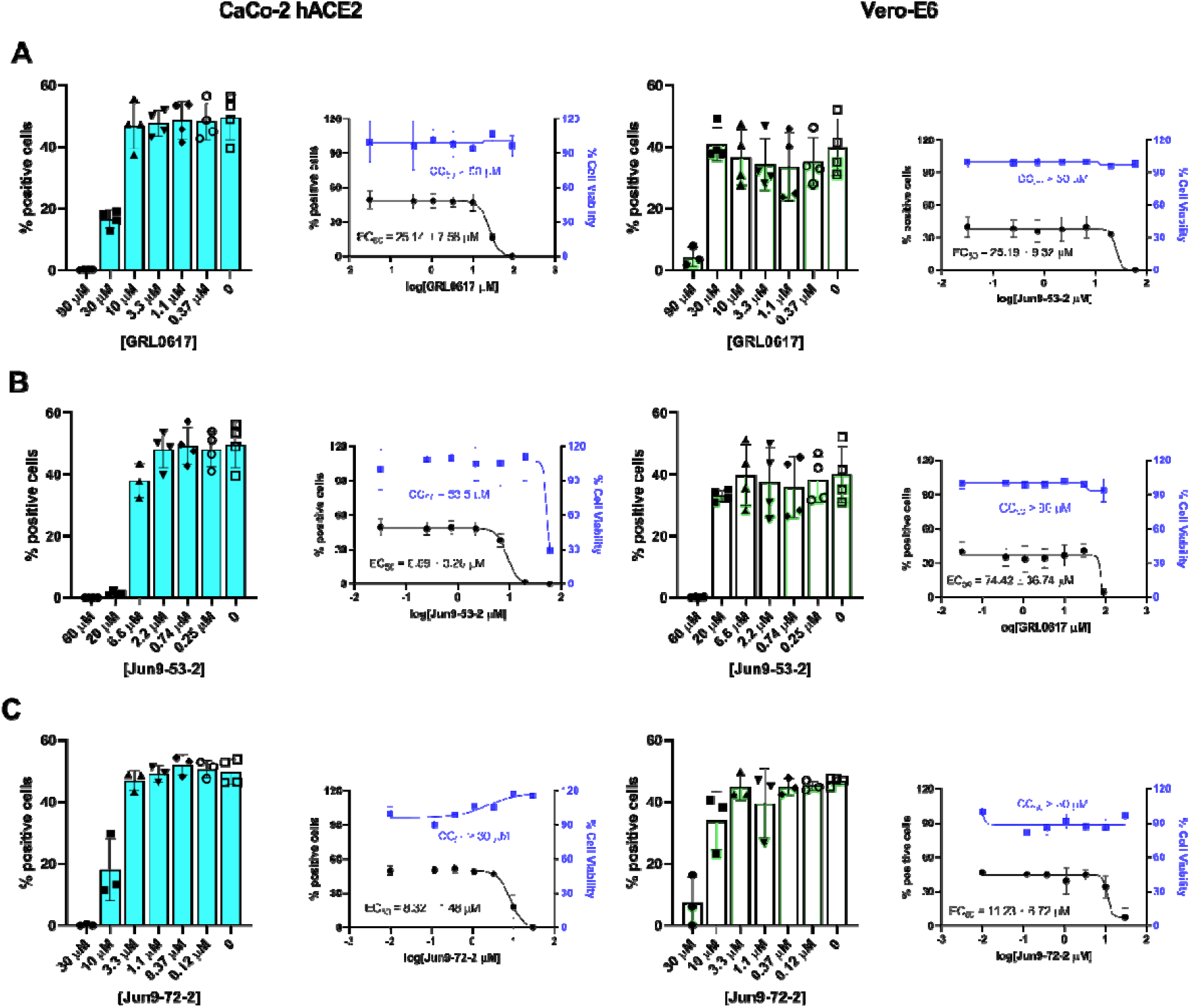
Cellular antiviral activity of Jun9-53-2 and Jun9-72-2 against SARS-CoV-2 in Caco-2 hACE2 and Vero E6 cells. (**A**) Antiviral activity of GRL0617. (**B**) Antiviral activity of Jun9-53-2. (**C**) Antiviral activity of Jun9-72-2.

### X-ray crystal structures of SARS-CoV-2 PL^pro^ in complex with GRL0617

The complex structure of SARS-CoV-2 PL^pro^ with GRL0617 was determined at 2.50 Å resolution, providing insight to its mechanism of inhibition. There are two monomers per asymmetric unit in the P2_1_ space group. Unambiguous electron density reveals GRL0617 binds to the S3-S4 subpockets of the PL^pro^ active site in a noncovalent manner (Fig. 6A). The naphthalene ring is positioned in the S4 site, where it forms hydrophobic interactions with Pro247 and Pro248. Upon ligand binding, Tyr268 flips towards the central domain of GRL0617 (Fig. 6B) to π-stack with the naphthalene and benzene rings. At the 1-napthalene position is a carbon stereocenter that is substituted with an (R) methyl group that sticks directly into the core of the S4 subpocket. This stereocenter and the methyl group are nearly superimposable with the γ and δ_1_ carbons of the P4 leucine for the ISG15 substrate in the previously determined complex structure (Fig. 6C), demonstrating the non-polar features of the S4 site as well as the complementarity of the methyl moiety with the core of this subpocket.^23^ The carboxamide nitrogen of GRL0617 serves as a hydrogen bond acceptor for the sidechain of Asp164, while the oxygen accepts a hydrogen bond from the mainchain amide of Gln269. The disubstituted benzene spans the central substrate channel, partially occupying the P5-P3 substrate mainchain binding site, where it forms π-π interactions with the sidechains of Tyr268/Gln269, and the backbone amides of Gly163/Asp164. The ortho-methyl group projects towards the catalytic core forming hydrophobic interactions with the S2 site, forcing Leu162 slightly outwards compared with the apo structure, while the meta-nitrogen orients towards the S5 site, causing Gln269 to swing inwards to accept a hydrogen bond. The conformational changes of multiple active site residues upon GRL0617 binding highlights the flexibility of the substrate recognition sites, which corresponds to the broad substrate profile of SARS-CoV-2 PL^pro^ and can be leveraged in future inhibitor design.

**Fig 6.**
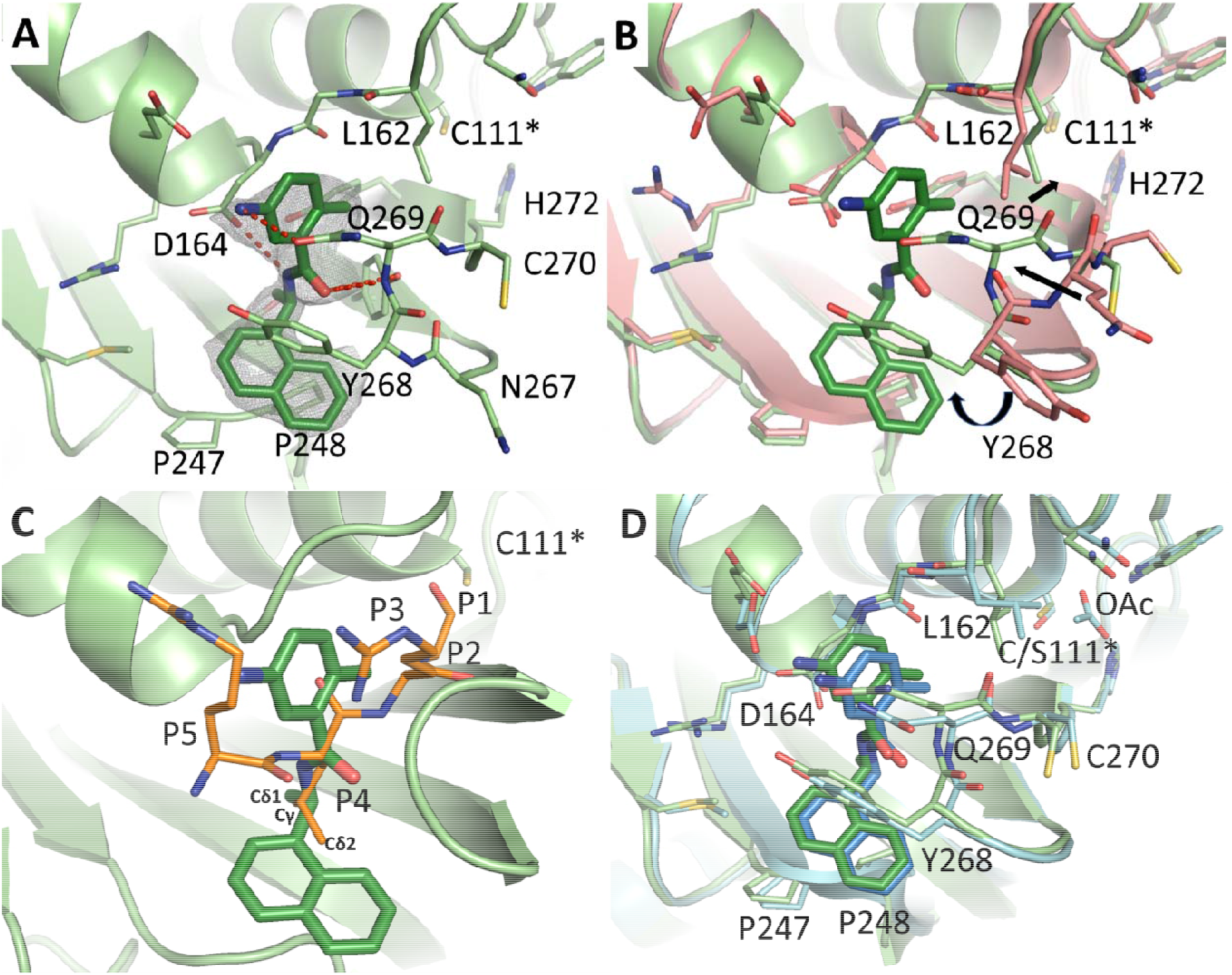
Complex structure of SARS-CoV-2 PL^pro^ with inhibitor GRL0617. The protein and ligand of the SARS-CoV-2 complex are colored in light green and dark green respectively. C111* indicates the catalytic cysteine. (**A**) Binding mode of GRL0617 with unbiased F_o_-F_c_ map, shown in grey, contoured at 2_σ_. Hydrogen bonds are shown as red dashed lines. (**B**) Superimposition with apo SARS-CoV-2 PL^pro^ (salmon). Significant rearrangement is observed in the loop comprising residues Asn267-Cys270 upon GRL0617 binding. These movements are indicated with arrows. (**C**) Superimposition with the terminal 5 residues of the ubiquitin like protein ISG15 substrate (orange) from the complex with SARS-COV-2 PL^pro^ (PDB ID 6XA9, showing only the ligand). The atoms of the Leu side chain at the P4 position are labeled. (**D**) Superimposition with the catalytic mutant C111S in complex with GRL0617 (PDB ID 7JIR, light blue and dark blue). OAc indicates an acetate molecule in the C111S mutant active site.

One of the unique aspects of GRL0167 is that it does not interact with the catalytic core, but instead binds to a distal portion of the active site. Other research groups have determined complex structures of PL^pro^ with GRL0617 with its catalytic cysteine, Cys 111, mutated to a serine, presumably to increase its propensity to crystallize (PDB IDs: 7JIR (2.1 Å) and 7CJM (3.2 Å)).^24,25^ When the three structures are compared, the GRL0167 adopts a nearly identical pose. Minor differences in the side chain conformations of Glu167 and Gln269 are observed. However, there is a significant difference in the pose of Leu162 between the WT and the C111S mutants (Fig. 6D). In our WT structure, Leu162 inserts into the core of the protein, where it maintains an interatomic distance of 3.4 Å with the catalytic cysteine. In contrast, Leu162 of both C111S structures flip outwards, towards the solvent. In the higher resolution structure (PDB ID 7JIR), an acetate from the crystallization condition is modelled in the active site. When superimposed with our WT structure, this acetate clashes with Cys111 (closest distance 2.5 Å) and Leu162 (3.0 Å). In the lower-resolution C111S mutant complexed with GRL0617 (PDB ID 7CJM), no acetate is modelled, but Leu162 adopts the same conformation as the higher-resolution C111S structure (PDB ID 7JIR). Further inspection of the 2F_o_-F_c_ map of 7JCM reveals there is unmodeled density corresponding to the acetate from PDB ID 7JIR. Interestingly, this experiment did not use acetate in their crystallization condition. Therefore, the density in the catalytic core of both C111S structures likely corresponds to a species of unknown identity that preferentially interacts with a serine residue.

## CONCLUSION

Given the devastating impact of the COVID-19 pandemic, the SARS-CoV outbreak in 2003 was a dire warning that was gravely overlooked in retrospect. Looking forward, it is imperative that therapeutics are developed that are not only effective against SARS-CoV-2, but against future strains of similar coronaviruses. PL^pro^ is a high profile drug target, partly because it is highly conserved between SARS-CoV and SARS-CoV-2, sharing 83% sequence similarity. Furthermore, inhibitors like GRL0617 are equally effective against both viruses, with a Ki of 0.49 µM and 0.57 µM, against SARS-CoV PL^pro^ and SARS-CoV-2 PL^pro^.^26^ Likewise, all critical active site residues that interact with GRL0617 are conserved, consequently, the binding poses are nearly identical (fig. S4).^26^ These similarities would indicate that PL^pro^ inhibitors might retain their activity against future strains of SARS coronaviruses.

Previous attempts to discover SARS-CoV-2 PL^pro^ inhibitors through HTS by others have failed to identify hits with improved enzymatic inhibition and cellular antiviral activity.^8,9^ Structural analogs of GRL0617 were also designed and synthesized, however, none of them showed improved enzymatic inhibition.^15^ Part of the reason for the difficulty in targeting SARS-CoV-2 PL^pro^ is the lack of S1 and S2 pockets, which leaves only S3 and S4 pockets for inhibitor binding. Among the reported SARS-CoV or SARS-CoV-2 PL^pro^ inhibitors, GRL0617 is one of the most potent compound. However, it had weak antiviral activity (EC_50_ > 20 µM). In this study, we attempted to identify more potent SARS-CoV-2 PL^pro^ inhibitors through a HTS. Based on two promising hits Jun9-13-7 and Jun9-13-9, a library of analogs was designed and synthesized, among which several compounds had submicromolar IC_50_ values in the FRET-based enzymatic assay. To alleviate the burden of relying on BSL-3 facility to test the antiviral activity of PL^pro^ inhibitors, we developed the cell-based FlipGFP PL^pro^ assay, which can be used to quantify the intracellular enzymatic inhibition of PL^pro^ in a BSL-2 lab. The FlipGFP PL^pro^ assay is a close mimetic of the virus-infected cell in which PL^pro^ cleaves its substrate in the native intracellular reducing environment. The advantage of the FlipGFP PL^pro^ assay overt the standard FRET-based enzymatic assay is that it can rule out compounds that are either cytotoxic or membrane impermeable or non-specifically modify the catalytic cysteine through oxidation or alkylation. GRL0617 showed an EC_50_ of 9.29 µM in the FlipGFP PL^pro^ assay, while the SARS-CoV-2 M^pro^ inhibitor GC376 was not active (EC_50_ > 60 µM), suggesting this assay was suitable for testing PL^pro^ inhibitors. Among the ten PL^pro^ inhibitors tested in the FlipGFP PL^pro^ assay, five compounds Jun9-53-2, Jun9-72-2, Jun9-75-4, Jun9-85-1, and Jun9-87-1 had EC_50_ values less than 10 µM. Next, two prioritized hits Jun9-53-2 and Jun9-72-2 were advanced to the cellular antiviral assay against SARS-CoV-2. Both compounds showed improved antiviral activity than GRL0617 and the EC_50_ values were consistent between the FlipGFP PL^pro^ assay and the cellular antiviral assay.

We also solved the X-ray crystal structure of the wild-type SARS-CoV-2 PL^pro^ in complex with GRL0617. Binding of GRL0617 to SARS-CoV-2 induced a conformational change in the BL2 loop to the more closed conformation. In contrast, a larger inhibitor VIR251 stabilizes the BL2 loop in the open conformation.^5^ The intrinsic flexibility of the BL2 loop implies that structurally diverse inhibitors might be able to fit in the S3-S4 pockets.

In conclusion, the SARS-CoV-2 PL^pro^ inhibitors developed in this study represent promising hits for further development as SARS-CoV-2 antivirals, and the FlipGFP PL^pro^ assay is a suitable surrogate for testing the cellular activity of PL^pro^ inhibitors in the BSL-2 setting.

## MATERIALS AND METHODS

### Cell lines and viruses

VERO E6 cells (ATCC, CRL-1586) were cultured in Dulbecco’s modified Eagle’s medium (DMEM), supplemented with 5% heat inactivated FBS in a 37°C incubator with 5% CO_2_.

SARS-CoV-2, isolate USA-WA1/2020 (NR-52281), was obtained through BEI Resources and propagated once on VERO E6 cells before it was used for this study. Studies involving the SARS-CoV-2 were performed at the UTHSCSA biosafety level-3 laboratory by personnel wearing powered air purifying respirators.

### Protein expression and purification

Detailed expression and purification of C-terminal His tagged SARS-CoV-2 PL^Pro^ (PL^pro^-His) were described in our previous publication.^4^ Briefly, SARS-CoV-2 papain-like protease (PL^pro^) gene (ORF 1ab 1564−1876) from strain BetaCoV/ Wuhan/WIV04/2019 with *E. coli* codon optimization in the pET28b(+) vector was ordered from GenScript. The pET28b(+) plasmid was transformed into BL21(DE3) cells, and protein expression was induced with 0.5 mM IPTG when OD600 was around 0.8 for 24 h at 18 °C. Then cells were harvested and lysed, PL^pro^-His protein was purified with a single Ni-NTA resin column, eluted PL^pro^-His was dialyzed against 100-fold volume dialysis buffer (50 mM Tris [pH 7.5], 150 mM NaCl, 2 mM DTT) in a 10 000 kDa molecular weight cutoff dialysis tubing.

The expression and purification of un-tagged SARS-CoV-2 PL^pro^ (PL^pro^) was carried out as follows: the SARS-CoV-2 PL^pro^ gene (ORF 1ab 1564−1876) was subcloned from pET28b(+) to pE-SUMO vector according to the manufacturer’s protocol (LifeSensors Inc., Malvern, PA). The forward primer with the Bsa I site is GCGGTCTCAAGGTGAAGTTCGCACCATCAAAGTTTTTACC; the reverse primer with Xba I site is GCGGTCTCTCTAGATTACTTGATGGTGGTGGTGTAGCTGTTCTC. SUMO tagged protein was expressed and purified as PL^pro^-His protein. SUMO-tag was removed by incubation with SUMO protease 1 at 4°C overnight, and the free SUMO tag was removed by application of another round of Ni-NTA resin. The purity of the protein was confirmed with SDS-PAGE gel.

The expression and purification of SARS-CoV-2 M^pro^ with unmodified N- and C-termini were reported in previous studies.^4^

### Peptide synthesis

The SARS-CoV-2 PL^pro^ FRET substrate Dabcyl-FTLRGG/APTKV(Edans) and the SARS-CoV-2 M^pro^ FRET substrate Dabcyl-KTSAVLQ/SGFRKME(Edans) were synthesized by solid-phase synthesis through iterative cycles of coupling and deprotection using the previously optimized procedure.^27^Ub-AMC and ISG15-AMC were purchased from Boston Biochem (catalog # U-550-050 and UL-553-050 respectively).

### Compound synthesis and characterization

Details for the synthesis procedure and characterization for compounds can be found in the supplementary information.

### Enzymatic assays

The high-throughput screening was carried out in 384-well format. 1 µl of 2 mM library compound was added to 50 µl of 200 nM PL^pro^-His protein in PL^pro^ reaction buffer (50 mM HEPES pH7.5, 5 mM DTT and 0.01% Triton X-100) and was incubated at 30 °C for 1 h. the reaction was initiated by adding 1 µl of 1 mM PL^pro^ FRET substrate. The end point fluorescence signal was measured after 3 h incubation at 30 °C with a Cytation 5 image reader with filters for excitation at 360/40 nm and emission at 460/40 nm. The final testing compound concentration is ∼ 40 µM and FRET substrate concentration is ∼ 20 µM; a control plate as in Fig.1 was included in every batch of screening.

The diversity compound library consisting of 50,240 compounds was purchased from Enamine (Cat # 781270).

For the measurements of *K*_m_/*V*_max_: with Peptide-Edans as a substrate, the final PL^pro^ protein concentration is 200 nM, and substrate concentration ranges from 0 to 200 µM; with Ub-AMC as a substrate, the final PL^pro^ protein concentration is 50 nM, and Ub-AMC concentration ranges from 0 to 40 µM; with ISG15-AMC as a substrate, the final PL^pro^ protein concentration is 2 nM, and ISG15-AMC concentration ranges from 0 to 15 µM. The reaction was monitored in a Cytation 5 image reader with filters for excitation at 360/40 nm and emission at 460/40 nm at 30 °C for 1 hr. The initial velocity of the enzymatic reaction was calculated from initial 10 min enzymatic reaction and was plotted against with substrate concentrations in Prism 8 with a Michaelis-Menton function.

For the IC_50_ measurement with FRET Peptide-Edans substrate: the reaction was carried out in 96-well format with 200 nM PL^pro^ protein as described previously^3,4^. For the IC_50_ measurements with Ub-AMC or ISG15-AMC substrate, the reaction was carried out in 384-well format. The final PL^pro^ protein concentration is 50 nM and substrate concentration is 2.5 µM when Ub-AMC is applied; The final PL^pro^ protein concentration is 2 nM and substrate concentration is 0.5 µM when ISG15-AMC is applied. For the Lineweaver-Burk plots of GRL0617, Jun9-13-7 and Jun9-13-9, assay was carried as follows: 50 µl of 400 nM PL^pro^ protein was added to 50 µl reaction buffer containing testing compound and various concentrations of FRET Peptide-Edans substrate to initiate the enzyme reaction. The initial velocity of the enzymatic reaction with and without testing compounds was calculated by linear regression for the first 10 min of the kinetic progress curve, the plotted against substrate concentrations in Prism 8 with Michaelis-Menten equation and linear regression of double reciprocal plot.

The main protease (M^pro^) enzymatic assays were carried out in M^pro^ reaction buffer containing 20 mM HEPES pH 6.5, 120 mM NaCl, 0.4 mM EDTA, 20% glycerol and 4 mM DTT as described previously.^3,4,19,20^

### Cell-based FlipGFP PL^pro^ assay

Plasmid pcDNA3-TEV-flipGFP-T2A-mCherry was ordered from Addgene (Cat # 124429). SARS CoV-2 PL^pro^ cleavage site LRGGAPTK or SARS CoV-2 M^pro^ cleavage site AVLQSGFR was introduced into pcDNA3-flipGFP-T2A-mCherry via overlapping PCRs to generate a fragment with SacI and HindIII sites at the ends. SARS CoV-2 M^pro^ and PL^pro^ expression plasmids pcDNA3.1 SARS2 M^pro^ and pcDNA3.1 SARS2 PL^pro^ was ordered from Genscript (Piscataway NJ) with codon optimization.

For transfection, 96-well Greiner plate (Cat # 655090) was seeded with 293T cells to overnight 70-90% confluency. 50 ng pcDNA3-flipGFP-T2A-mCherry plasmid and 50 ng protease expression plasmid pcDNA3.1 were used each well in the presence of transfection reagent TransIT-293 (Mirus). 3 hrs after transfection, 1 µl testing compound was added to each well at 100-fold dilution. Images were acquired 2 days after transfection with Cytation 5 imaging reader (Biotek) GFP and mCherry channels; and were analyzed with Gen5 3.10 software (Biotek). SARS CoV-2 PL^pro^ protease activity was calculated by the ration of GFP signal sum intensity over mCherry signal sum intensity. FlipGFP-PLP assay IC_50_ value was calculated by plotting GFP/mCherry signal over the applied compound concentration with a 4 parameters dose-response function in prism 8. The mCherry signal alone was utilized to determine the compound cytotoxicity.

### Differential scanning fluorimetry (DSF)

The thermal shift binding assay (TSA) was carried out using a Thermal Fisher QuantStudio™ 5 Real-Time PCR System as described previously.^3,4,19,20^ Briefly, 4 µM SARS-CoV-2 PL^pro^ protein (PL^pro^) in PL^pro^ reaction buffer was incubated with 40 µM of compounds at 30 °C for 30 min. 1X SYPRO orange dye was added and fluorescence of the well was monitored under a temperature gradient range from 20 °C to 90 °C with 0.05 °C/s incremental step. Measured T_m_ was plotted against compound concentration with one-site binding function in Prism 8.

### Native Mass Spectrometry

Before MS analysis, the protein was buffer exchanged using two Micro Bio-Spin columns (Bio-Rad) and diluted into 0.2M ammonium acetate to a concentration of 10 µM. Each drug was diluted with 100% ethanol to concentrations of 100, 200, and 400 μM. Imidazole, a charge reducing reagent, was added to each sample to stabilize the drug bound state at final concentration of 40 mM. For each sample, 0.5 μl ligand was added and dried down in each tube prior to the addition of 0.5 μL of 40 mM DTT, 0.5 μl of 400 mM imidazole, and 4 μl of protein for a final concentration of 4 mM DTT and 8 μM protein. Final ligand concentrations were either 10, 20, or 40 μM.

Native mass spectrometry (MS) was performed using a Q-Exactive HF quadrupole-Orbitrap mass spectrometer with the Ultra-High Mass Range research modifications (Thermo Fisher Scientific) as described in our previous publications^3,4^.

### Immunofluorescence assay

Antiviral immunofluorescence assay was carried out as previously described.^4,28^ Briefly, Vero E6 cells or Caca2-hACE2 cells in 96-well plates (Corning) were infected with SARS-CoV-2 (USA-WA1/2020 isolate) at a MOI of 0.1 in DMEM supplemented with 1% FBS. Immediately before the viral inoculation, the tested compounds in a three-fold dilution concentration series were also added to the wells in triplicate. The infection proceeded for 24 h without the removal of the viruses or the compounds. The staining and quantification procedures are described in our previous publications.^4^

### Crystallization and structure determination

SARS-CoV-2 PL^pro^-His (PL^pro^-His) protein was concentrated and loaded to a HiLoad 16/60 Superdex 75 size exclusion column (GE Healthcare) pre-equilibriated with 20 mM Tris pH 8.0 and 5 mM NaCl. Peak fractions were pooled and incubated with GRL0617 in a 1:1 molar ratio for 1 hour at room temperature, then concentrated to 8 mg/mL. PL^pro^ crystals were grown in a hanging-drop, vapor-diffusion apparatus by mixing 0.75 µl of 8 mg/mL PL^pro^-GRL0617 with 0.75 ul of well solution (30 % PEG 4,000, 0.2 M Li2SO4, and 0.1 M Tris pH 8.5). Crystals were transferred to a cryoprotectant solution containing 30 % PEG 4,000, 0.2 M Li_2_SO_4_, 0.1 M Tris pH 8.5 and 15 % glycerol, before being flash frozen in liquid nitrogen.

X-ray diffraction data for SARS-CoV-2 PL^pro^ + GRL0617 was collected on the SBC 19-BM beamline at the Advanced Photon Source (APS) in Argonne, IL, and processed with the HKL2000 software suite.^29^ The CCP4 versions of MOLREP was used for molecular replacement using a previously solved apo SARS-CoV-2 PL^pro^ structure, PDB ID: 6WZU as a reference model.^30,31^ Rigid and restrained refinements were performed using REFMAC and model building was performed with COOT.^32,33^ Protein structure figures were made using PyMOL (Schrödinger, LLC).

## DATA AVAILABILITY

The drug-bound complex structures for SARS-CoV-2 PL^pro^ with GRL0617 was deposited in the Protein Data Bank with accession numbers of 7JRN.

## ACKNOWLEDGEMENTS

This research was partially supported by the National Institutes of Health (NIH) (Grants AI147325 and AI147046) and the Arizona Biomedical Research Centre Young Investigator grant (ADHS18-198859) to J. W. J. A. T. and M. T. M. were funded by the National Institute of General Medical Sciences and National Institutes of Health (Grant R35 GM128624 to M. T. M.). We thank Michael Kemp for assistance with crystallization and X-ray diffraction data collection. We also thank the staff members of the Advanced Photon Source of Argonne National Laboratory, particularly those at the Structural Biology Center (SBC), with X-ray diffraction data collection. SBC-CAT is operated by UChicago Argonne, LLC, for the U.S. Department of Energy, Office of Biological and Environmental Research under contract DE-AC02-06CH11357. The SARS-CoV-2 experiments were supported by a COVID-19 pilot grant from UTHSCSA and NIH grant AI151638 to Y.X. SARS-Related Coronavirus 2, Isolate USA-WA1/2020 (NR-52281) was deposited by the Centers for Disease Control and Prevention and obtained through BEI Resources, NIAID, NIH.

## AUTHOR CONTRIBUTIONS

J. W. and C. M. conceived and designed the study; Z. X. and N. K. designed and synthesized the PL^pro^ inhibitors; C. M. expressed the PL^pro^ with the assistance of T. S.; C. M. performed the HTS, IC_50_ determination, thermal shift-binding assay, and enzymatic kinetic studies with the assistance of M. B.; C. M. developed the FlipGFP PL^pro^ assay. M. S. carried out M^pro^ crystallization and structure determination with the assistance of X. Z, and analyzed the data with Y. C.; J. T. performed the native mass spectrometry experiments with the guidance from M. M.; Y. H. performed the cytotoxicity assay; X. M. and F. Z. performed the SARS-CoV-2 immunofluorescence assay under the guidance of Y. X.; J. W. and Y. C. secured funding and supervised the study; J. W., Y.C., and M.S. wrote the manuscript with the input from others.

## ADDITIONAL INFORMATION

Supplementary information accompanies this paper at

