## Supplementary Information for "Discovery of SARS-CoV-2 papain-like protease inhibitors through a combination of high-throughput screening and FlipGFP-based reporter assay"

Table S1-S5

Fig. S1-S4

**Table S1: Enzymatic activities of SARS-CoV-2 PL^pro^-His and SARS-CoV-2 PL^pro^ in cleaving the FRET substrate.**

|  | SAR-CoV-2 PL^pro^-His | SARS-CoV-2 PL^pro^ |
| --- | --- | --- |
| k_cat_ (s^-1^) | 0.0245 | 0.0160 |
| K_m_ (µM^-1^) | 72.0 | 63.3 |
| k_cat_/K_m_ (M^-1^ s^-1^) | 340 | 255 |

**Table S2: Enzymatic activities of SARS-CoV-2 PL^pro^-His and SARS-CoV-2 PL^pro^ in cleaving the ubiquitin and ISG substrates.**

| SARS-CoV-2 PL^pro^-His | FRET substrate | Ub-AMC | ISG-AMC |
| --- | --- | --- | --- |
| k_cat_ (s^-1^) | 0.0245 | 0.035 | 1.293 |
| K_m_ (µM^-1^) | 72.0 | 32.7 | 7.75 |
| k_cat_/K_m_ (M^-1^ s^-1^) | 340 (1X) | 1070 (3X) | 1.67x10^5^ (490X) |

**Table S3: Validation of the HTS hits in the secondary FRET-based enzymatic assay and DSF binding assay.**

| Compounds | SARS CoV-2  IC_50_ (µM) | DSF  ΔTm (°C) | Compounds | SARS CoV-2  IC_50_ (µM) | DSF  ΔTm (°C) |
| --- | --- | --- | --- | --- | --- |
| GRL-0617 | 1.60 ± 0.13 | 3.52 ± 0.27 | Jun9-13-1 | 78.1 ± 28.1 | -0.45 ± 0 |
| Jun9-10-1 | 16.7 ± 2.48 | -2.96 ± 0.04 | Jun9-13-2 | >200 | -0.24 ± 0.35 |
| Jun9-10-2 | 98.8 ± 13.2 | 0.93 ± 0.28 | Jun9-13-3 | 18.2 ± 8.55 | 1.01 ± 0.07 |
| Jun9-10-3 | 19.6 ± 1.6 | 0.22 ± 0.00 | Jun9-13-4 | 144 ± 37 | 0.15 ± 0.36 |
| Jun9-10-4 | 16.2 ± 5.9 | 0.71 ± 0.07 | Jun9-13-5 | >200 | -- |
| Jun9-10-5 | 12.8 ± 2.3 | 0.15 ± 0.35 | Jun9-13-6 | >200 | 0.45 ± 0.07 |
| Jun9-10-6 | 23.7 ± 10.1 | 1.21 ± 0.42 | Jun9-13-7 | 7.29 ± 1.03 | 2.98 ± 0.09 |
| Jun9-10-7 | 17.4 ± 2.2 | 0.81 ± 0.07 | Jun9-13-8 | 71.8 ± 17.91 | 0.59 ± 0.07 |
| Jun9-10-8 | 20.5 ± 5.1 | 1.06 ± 0.28 | Jun9-13-9 | 6.67 ± 0.55 | 2.18 ± 0.29 |
| Jun9-10-9 | 38.7 ± 2.9 | 0.76 ± 0.14 | Jun9-13-1 | 78.1 ± 28.1 | -0.45 ± 0 |
|  |  |  | Jun9-13-2 | >200 | -0.24 ± 0.35 |

| Compound | FRET substrate  IC_50_ (µM) | Ub-AMC  IC_50_ (µM) | ISG-AMC  IC_50_ (µM) |
| --- | --- | --- | --- |
| **GRL0617** | 1.68 ± 0.22 | 0.88 ± 0.59 | 1.68 ± 0.39 |
| **Jun9-13-9** | 6.67 ± 0.55 | 4.93 ± 0.76 | 8.19 ± 2.60 |
| **Jun9-13-7** | 7.29 ± 1.03 | 6.58 ± 1.68 | 12.5 ± 5.1 |

**Table S4: Inhibitory activity of Jun9-13-9 and Jun9-13-7 against SARS-CoV-2 PL^pro^ using different substrates.**

**Table S5. Crystallographic statistics**

| **Data Collection** | **PDB ID 7JRN** |
| --- | --- |
| Inhibitor | GRL0617 |
| Space Group | P 1 2_1_ 1 |
| Cell Dimension |  |
| a, b, c (Å) | 46.73, 144.87, 60.15 |
| α, β, γ (°) | 90.00, 99.27, 90.00 |
| Resolution (Å) | 50.00 – 2.50 |
|  | (2.54 – 2.50) |
| R_merge_ | 0.128 (0.384) |
| <I>/σ<I> | 9.04 (2.02) |
| Completeness (%) | 86.2 (66.6) |
| Redundancy | 7.2 (6.8) |
| **Refinement** |  |
| Resolution (Å) | 39.64 – 2.48 |
|  | (2.56 – 2.48) |
| No. reflections/free | 23798 / 1175 |
| R_work_/R_free_ | 0.25 / 0.29 |
| No. Atoms | 5160 |
| Protein | 4998 |
| Ligand/Ion | 73 |
| Water | 89 |
| B-Factors (Å^2^) | 37.15 |
| Protein | 37.18 |
| Ligand/Ion | 45.81 |
| Solvent | 28.43 |
| RMS Deviations |  |
| Bond Lengths (Å) | 0.014 |
| Bond Angles (°) | 1.69 |
| Ramachandran Favored (%) | 94.41 |
| Ramachandran Allowed (%) | 5.27 |
| Ramachandran Outliers (%) | 0.32 |

* Values in parentheses refer to the last resolution shell.

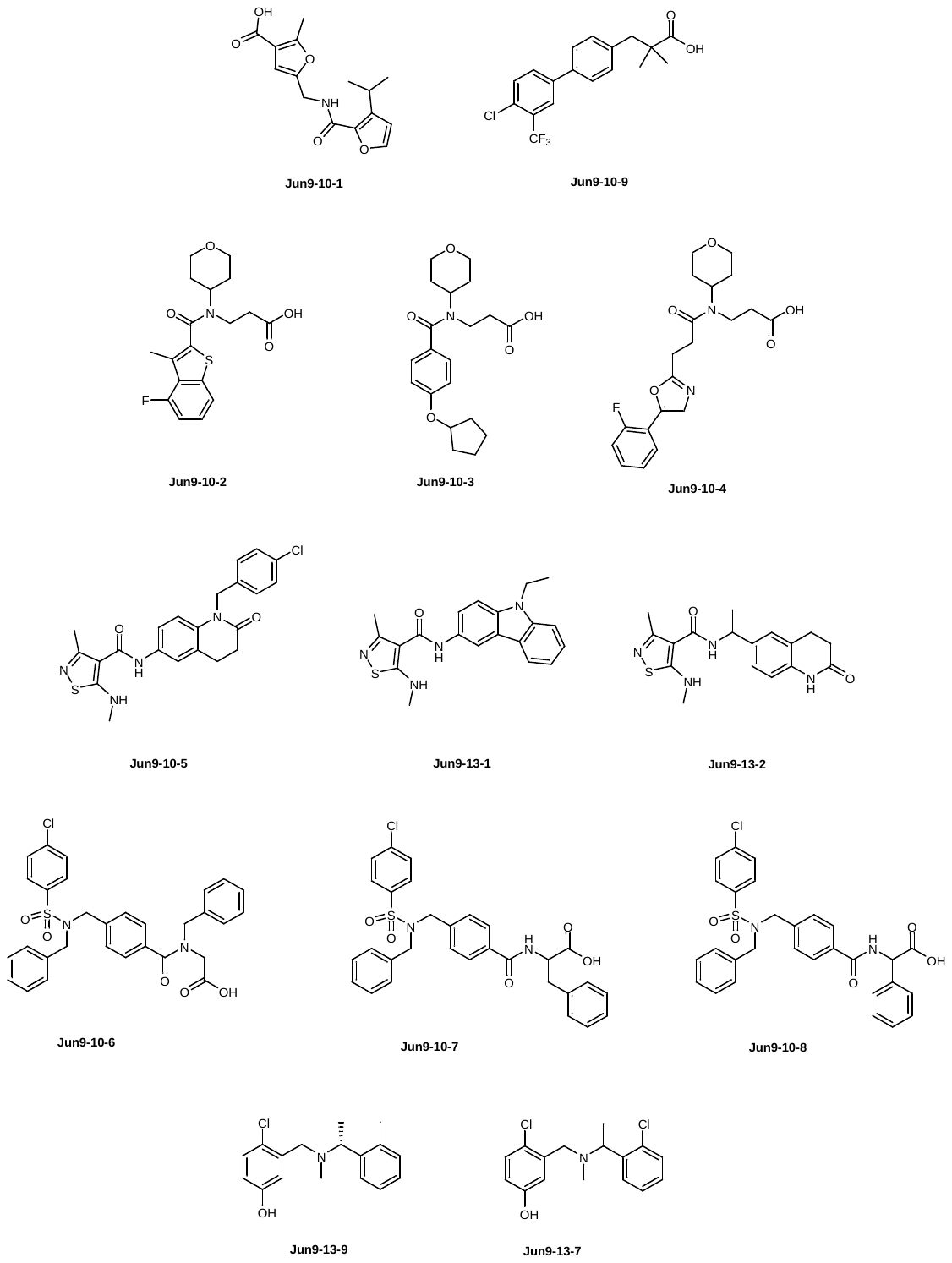

**Fig. S1. Chemical structures of the hits from HTS of the Enamine 50K diversity library against SARS-CoV-2 PL^pro^.**

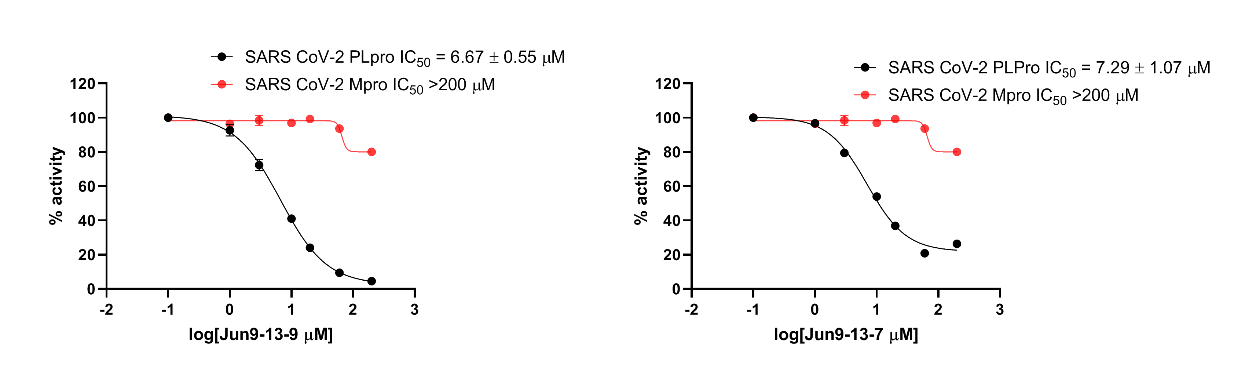

**Fig. S2. Selectivity profiling of the Jun9-13-9 and Jun9-13-7 against SARS-CoV-2 M^pro^.**

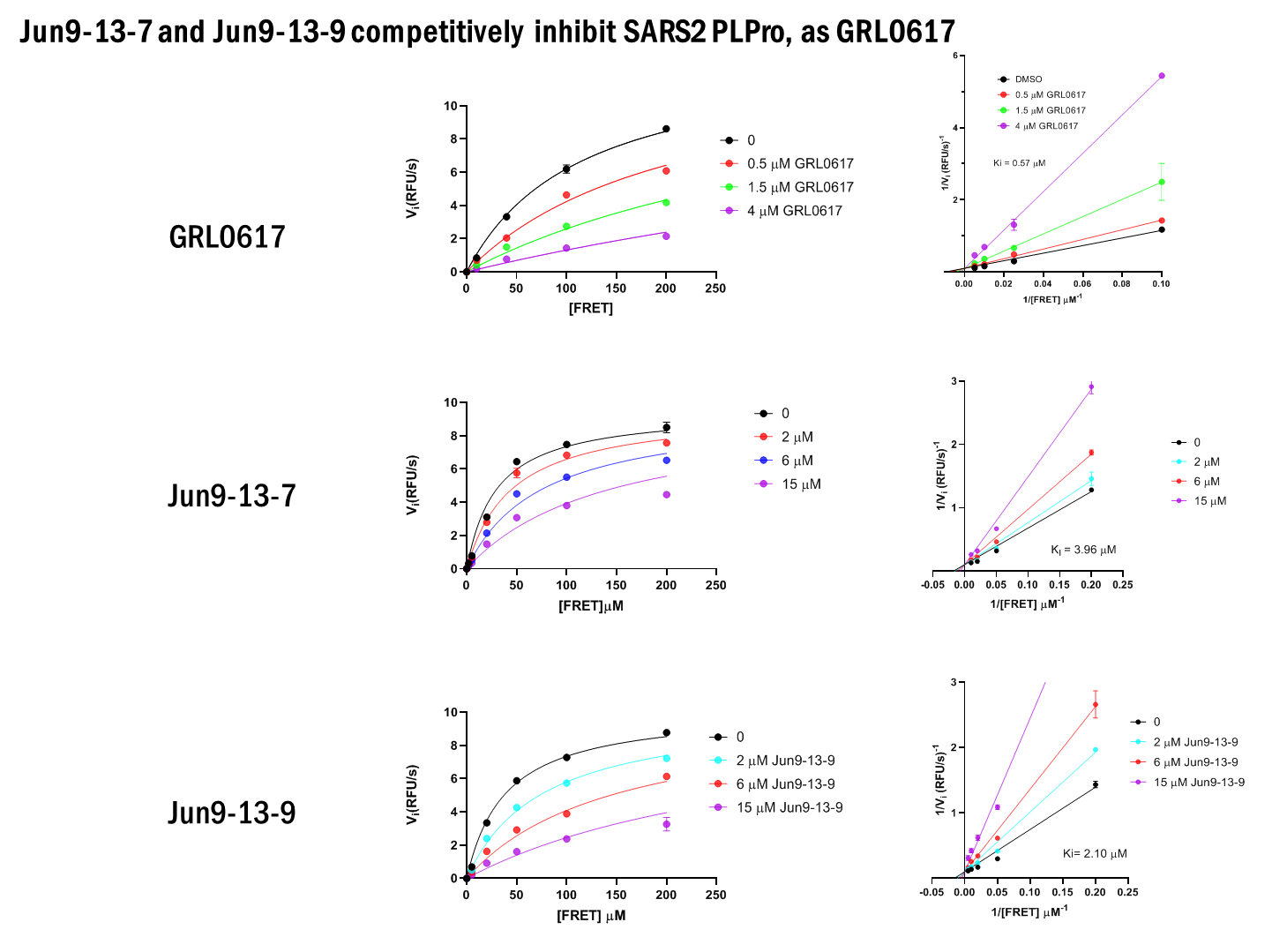

**Fig. S3. Enzymatic kinetic studies and Lineweaver-Burk plots of Jun9-13-7 and Jun9-13-9 in inhibiting SARS-CoV-2 PL^pro^.**

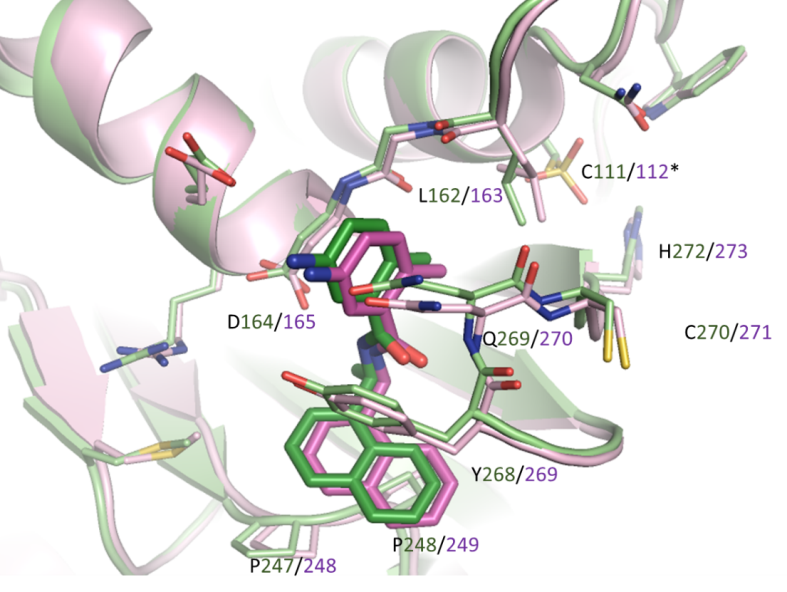

**Fig. S4. Superimposition of SARS-CoV PL^pro^ (light green) and SARS-CoV-PL^pro^ (lavender) with GRL0617 (dark green / purple) (PDB ID 3E9S).**

**Chemistry.** Chemicals were ordered from commercial sources and were used without further purification. All final compounds were purified by flash column chromatography. ^1^H and ^13^C NMR spectra were recorded on a Bruker-400 NMR spectrometer. Chemical shifts are reported in parts per million referenced with respect to residual solvent DMSO-d6) 2.50 ppm and from internal standard tetramethylsilane (TMS) 0.00 ppm. The following abbreviations were used in reporting spectra: s, singlet; d, doublet; t, triplet; q, quartet; m, multiplet; dd, doublet of doublets. All reactions were carried out under N_2_ atmosphere unless otherwise stated. HPLC-grade solvents were used for all reactions. Flash column chromatography was performed using silica gel (230-400 mesh, Merck). Low-resolution mass spectra were obtained using an ESI technique on a 3200 Q Trap LC/MS/MS system (Applied Biosystems). The purity was assessed by using Shimadzu LC-MS with Waters XTerra MS C-18 column (part #186000538), 50 × 2.1 mm, at a flow rate of 0.3 mL/min; λ = 250 and 220 nm; mobile phase A, 0.1% formic acid in H_2_O, and mobile phase B’, 0.1% formic in 60% isopropanol, 30% CH_3_CN and 9.9% H_2_O.

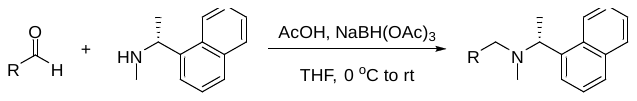

**General Synthesis Procedure**. The PL^pro^ inhibitors were synthesized as the following procedure unless otherwise noted. To the solution of (*R*)-N-methyl-1-(naphthalen-1-yl)ethan-1-amine (1 mmol) in anhydrous THF (10 mL) was added the corresponding aldehyde (1 mmol), followed by AcOH (1.05 mmol). Then, the mixture was cooled with ice-water batch and was added NaBH(OAc)_3_ (3 mmol). The reaction was warmed to room temperature and stirred overnight. After removing THF completely, the residue was added 1 N HCl (2 mL) and the mixture was washed with hexane/CH_2_Cl_2_ (hexane/CH_2_Cl_2_ = 2:1) and the organic layer was discarded. Then, the aqueous layer was adjusted pH to slightly basic with 2.5 N NaOH and the mixture was extracted with CH_2_Cl_2_/MeOH (CH_2_Cl_2_/MeOH = 15:1). The combined organic layer was separated, dried over anhydrous Na_2_SO_4_, filtered and concentrated under reduced pressure. The residue was purified by silica gel flash column chromatography (CH_2_Cl_2_ to CH_2_Cl_2_/MeOH = 10:1) to afford the target product.

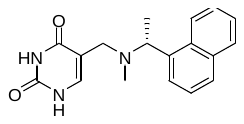

Jun9-81-1. Yield: 75%. ^1^H NMR (400 MHz, DMSO-d6) δ 11.03 (s, 1H), 10.66 (s, 1H), 8.45-8.40 (m, 1H), 7.93-7.88 (m, 1H), 7.80 (d, *J* = 8.0 Hz, 1H), 7.55 (d, *J* = 6.4 Hz, 1H), 7.52-7.43 (m, 3H), 7.10 (s, 1H), 4.48 (q, *J* = 6.4 Hz, 1H), 3.30 (d, *J* = 14.0 Hz, 1H), 3.17 (d, *J* = 14.0 Hz, 1H), 2.06 (s, 3H), 1.45 (d, *J* = 6.4 Hz, 3H). ^13^C NMR (100 MHz, DMSO-d6) δ 164.34, 151.19, 139.73, 139.22, 133.62, 131.48, 128.39, 127.34, 125.52, 125.39, 125.16, 124.71, 124.16, 109.52, 58.83, 49.39, 37.03, 14.47. C_18_H_20_N_3_O_2_ ESI-MS: m/z (M + H^+^): 310.2 (calculated), 310.2 (found).

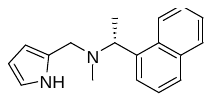

Jun9-67-2. Yield: 82%. ^1^H NMR (400 MHz, DMSO-d6) δ 10.57 (s, 1H), 8.34-8.22 (m, 1H), 7.96-7.85 (m, 1H), 7.80 (d, *J* = 8.0 Hz, 1H), 7.63 (d, *J* = 7.2 Hz, 1H), 7.54-7.43 (m, 3H), 6.67 (dd, *J* = 4.0, 2.4 Hz, 1H), 5.96 (dd, *J* = 5.2, 2.4 Hz, 1H), 5.88 (s, 1H), 4.36 (q, *J* = 6.4 Hz, 1H), 3.60 (d, *J* = 13.6 Hz, 1H), 3.50 (d, *J* = 13.6 Hz, 1H), 2.06 (s, 3H), 1.45 (d, *J* = 6.4 Hz, 3H). ^13^C NMR (100 MHz, DMSO-d6) δ 140.26, 133.62, 131.33, 128.68, 128.40, 127.13, 125.60, 125.33, 125.29, 124.41, 124.32, 116.85, 107.17, 106.90, 57.86, 51.23, 37.73, 16.01. C_18_H_21_N_2_ ESI-MS: m/z (M + H^+^): 265.2 (calculated), 265.2 (found).

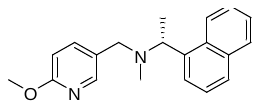

Jun9-68-1. Yield: 81%. ^1^H NMR (400 MHz, DMSO-d6) δ 8.42 (d, *J* = 8.4 Hz, 1H), 7.99-7.88 (m, 2H), 7.82 (d, *J* = 8.0 Hz, 1H), 7.66-7.39 (m, 5H), 6.71 (d, *J* = 8.4 Hz, 1H), 4.47 (q, *J* = 6.4 Hz, 1H), 3.80 (s, 3H), 3.52 (d, *J* = 13.2 Hz, 1H), 3.36 (d, *J* = 13.2 Hz, 1H), 2.06 (s, 3H), 1.49 (d, *J* = 6.4 Hz, 3H). ^13^C NMR (100 MHz, DMSO-d6) δ 162.71, 146.35, 139.81, 139.45, 133.70, 131.39, 128.51, 127.97, 127.43, 125.57, 125.46, 125.23, 124.56, 124.36, 110.06, 59.22, 54.46, 52.97, 37.26, 14.99. C_20_H_23_N_2_O ESI-MS: m/z (M + H^+^): 307.2 (calculated), 307.2 (found).

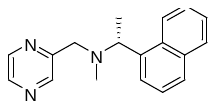

Jun9-68-4. Yield: 73%. ^1^H NMR (400 MHz, DMSO-d6) δ 8.55-8.38 (m, 4H), 7.95-7.88 (m, 1H), 7.83 (d, *J* = 8.0 Hz, 1H), 7.63 (d, *J* = 7.2 Hz, 1H), 7.59-7.46 (m, 3H), 4.58 (q, *J* = 6.4 Hz, 1H), 3.80 (d, *J* = 14.4 Hz, 1H), 3.68 (d, *J* = 14.4 Hz, 1H), 2.17 (s, 3H), 1.52 (d, *J* = 6.4 Hz, 3H). ^13^C NMR (100 MHz, DMSO-d6) δ 155.40, 144.54, 143.56, 142.86, 139.48, 133.67, 131.37, 128.51, 127.52, 125.63, 125.47, 125.22, 124.49, 124.38, 59.17, 57.47, 38.24, 14.99. C_18_H_20_N_3_ ESI-MS: m/z (M + H^+^): 278.2 (calculated), 278.2 (found).

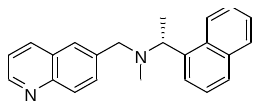

Jun9-68-3. Yield: 80%. ^1^H NMR (400 MHz, DMSO-d6) δ 8.84 (dd, *J* = 4.0, 1.6 Hz, 1H), 8.49 (d, *J* = 8.4 Hz, 1H), 8.25 (d, *J* = 7.6 Hz, 1H), 7.92 (t, *J* = 8.4 Hz, 2H), 7.83 (d, *J* = 8.0 Hz, 1H), 7.77 (s, 1H), 7.67 (d, *J* = 7.2 Hz, 1H), 7.63-7.44 (m, 5H), 4.54 (q, *J* = 6.4 Hz, 1H), 3.78 (d, *J* = 13.6 Hz, 1H), 3.62 (d, *J* = 13.6 Hz, 1H), 2.12 (s, 3H), 1.54 (d, *J* = 6.4 Hz, 3H). ^13^C NMR (100 MHz, DMSO-d6) δ 149.96, 147.12, 139.87, 138.27, 135.58, 133.69, 131.41, 130.23, 128.72, 128.51, 127.59, 127.41, 126.69, 125.58, 125.47, 125.25, 124.57, 124.35, 121.42, 59.35, 57.81, 37.77, 15.36. C_23_H_23_N_2_ ESI-MS: m/z (M + H^+^): 327.2 (calculated), 327.2 (found).

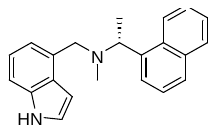

Jun9-75-4. Yield: 56%. ^1^H NMR (400 MHz, DMSO-d6) δ 11.01 (s, 1H), 8.58-8.37 (m, 1H), 7.91 (dd, *J* = 6.4, 2.8 Hz, 1H), 7.82 (d, *J* = 8.0 Hz, 1H), 7.68 (d, *J* = 7.2 Hz, 1H), 7.53-7.45 (m, 3H), 7.27 (d, *J* = 8.0 Hz, 1H), 7.23 (t, *J* = 2.8 Hz, 1H), 7.01 (t, *J* = 7.6 Hz, 1H), 6.94 (d, *J* = 7.2 Hz, 1H), 6.37 (s, 1H), 4.50 (q, *J* = 6.4 Hz, 1H), 3.78 (s, 2H), 2.10 (s, 3H), 1.56 (d, *J* = 6.4 Hz, 3H). ^13^C NMR (100 MHz, DMSO-d6) δ 140.30, 135.83, 133.64, 131.48, 130.70, 128.39, 127.26, 127.24, 125.44, 125.37, 125.22, 124.70, 124.52, 124.33, 120.63, 118.75, 110.00, 99.71, 59.45, 56.27, 38.15, 15.32. C_22_H_23_N_2_ ESI-MS: m/z (M + H^+^): 315.2 (calculated), 315.2 (found).

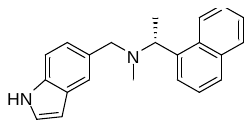

Jun9-80-4. Yield: 75%. ^1^H NMR (400 MHz, DMSO-d6) δ 10.99 (s, 1H), 8.47 (d, *J* = 8.0 Hz, 1H), 8.01-7.88 (m, 1H), 7.82 (d, *J* = 8.0 Hz, 1H), 7.66 (d, *J* = 6.8 Hz, 1H), 7.60-7.46 (m, 3H), 7.39 (s, 1H), 7.31-7.27 (m, 2H), 6.97 (dd, *J* = 8.4, 1.2 Hz, 1H), 6.42-6.28 (m, 1H), 4.44 (q, *J* = 6.4 Hz, 1H), 3.67 (d, *J* = 12.8 Hz, 1H), 3.47 (d, *J* = 12.8 Hz, 1H), 2.07 (s, 3H), 1.51 (d, *J* = 6.4 Hz, 3H). ^13^C NMR (100 MHz, DMSO-d6) δ 140.36, 135.04, 133.68, 131.44, 129.72, 128.45, 127.45, 127.23 , 125.44, 125.40, 125.25, 124.70, 124.32, 122.07, 119.87, 110.95, 100.78, 59.21, 58.92, 37.45, 15.45. C_22_H_23_N_2_ ESI-MS: m/z (M + H^+^): 315.2 (calculated), 315.2 (found).

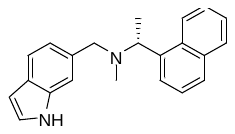

Jun9-68-2. Yield: 81%. ^1^H NMR (400 MHz, DMSO-d6) δ 10.96 (s, 1H), 8.48 (d, *J* = 8.0 Hz, 1H), 8.06-7.88 (m, 1H), 7.82 (d, *J* = 8.0 Hz, 1H), 7.66 (d, *J* = 7.2 Hz, 1H), 7.59-7.46 (m, 3H), 7.42 (d, *J* = 8.0 Hz, 1H), 7.29 (s, 1H), 7.28-7.25 (m, 1H), 6.89 (dd, *J* = 8.0, 1.2 Hz, 1H), 6.45-6.28 (m, 1H), 4.46 (d, *J* = 6.4 Hz, 1H), 3.71 (d, *J* = 13.2 Hz, 1H), 3.51 (d, *J* = 13.2 Hz, 1H), 2.09 (s, 3H), 1.52 (d, *J* = 6.4 Hz, 3H). ^13^C NMR (100 MHz, DMSO-d6) δ 140.29, 135.96, 133.69, 132.39, 131.45, 128.46, 127.27, 126.60, 125.50, 125.42, 125.24, 124.94, 124.66, 124.29, 119.95, 119.51, 111.14, 100.81, 59.29, 58.95, 37.52, 15.44. C_22_H_23_N_2_ ESI-MS: m/z (M + H^+^): 315.2 (calculated), 315.2 (found).

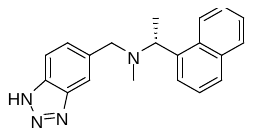

Jun9-84-2. Yield: 78%. ^1^H NMR (400 MHz, DMSO-d6) δ 8.47 (d, *J* = 8.4 Hz, 1H), 7.93 (d, *J* = 7.6 Hz, 1H), 7.83 (d, *J* = 8.0 Hz, 1H), 7.79 (d, *J* = 8.4 Hz, 1H), 7.69 (s, 1H), 7.65 (d, *J* = 7.2 Hz, 1H), 7.61-7.47 (m, 3H), 7.27 (d, *J* = 8.4 Hz, 1H), 4.54 (q, *J* = 6.4 Hz, 1H), 3.76 (d, *J* = 13.6 Hz, 1H), 3.60 (d, *J* = 13.6 Hz, 1H), 2.11 (s, 3H), 1.53 (d, *J* = 6.4 Hz, 3H). ^13^C NMR (100 MHz, DMSO-d6) δ 139.85, 133.70, 131.44, 128.52, 127.43, 125.58, 125.48, 125.23, 124.58, 124.33, 59.29, 57.85, 37.59, 15.04. C_20_H_21_N_4_ ESI-MS: m/z (M + H^+^): 317.2 (calculated), 317.2 (found).

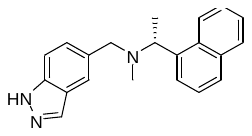

Jun9-85-1. Yield: 76%. ^1^H NMR (400 MHz, DMSO-d6) δ 8.48 (d, *J* = 8.4 Hz, 1H), 7.98 (s, 1H), 7.95-7.89 (m, 1H), 7.83 (d, *J* = 8.0 Hz, 1H), 7.67-7.45 (m, 5H), 7.40 (s, 1H), 6.97 (d, *J* = 8.4 Hz, 1H), 4.51 (q, *J* = 6.4 Hz, 1H), 3.74 (d, *J* = 13.2 Hz, 1H), 3.56 (d, *J* = 13.2 Hz, 1H), 2.11 (s, 3H), 1.52 (d, *J* = 6.4 Hz, 3H). ^13^C NMR (100 MHz, DMSO-d6) δ 140.18, 140.01, 138.03, 133.69, 133.11, 131.43, 128.49, 127.36, 125.56, 125.46, 125.23, 124.58, 124.29, 121.96, 121.33, 120.00, 109.10, 59.35, 58.42, 37.62, 15.27. C_21_H_22_N_3_ ESI-MS: m/z (M + H^+^): 316.2 (calculated), 316.2 (found).

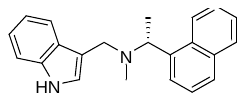

Jun9-84-3. Yield: 67%. ^1^H NMR (400 MHz, DMSO-d6) δ 10.86 (s, 1H), 8.40 (d, *J* = 8.4 Hz, 1H), 7.91 (d, *J* = 8.0 Hz, 1H), 7.82 (d, *J* = 8.0 Hz, 1H), 7.65 (d, *J* = 7.2 Hz, 1H), 7.55-7.28 (m, 5H), 7.20 (d, *J* = 2.0 Hz, 1H), 7.04 (t, *J* = 7.6 Hz, 1H), 6.87 (t, *J* = 7.6 Hz, 1H), 4.46 (q, *J* = 6.4 Hz, 1H), 3.73 (d, *J* = 13.2 Hz, 1H), 3.67 (d, *J* = 13.2 Hz, 1H), 2.10 (s, 3H), 1.53 (d, *J* = 6.4 Hz, 3H). ^13^C NMR (100 MHz, DMSO-d6) δ 140.82, 136.80, 134.05, 131.87, 128.74, 127.71, 127.58, 125.72, 125.60, 125.22, 124.80, 124.75, 121.22, 119.48, 118.54, 112.41, 111.64, 59.03, 49.76, 38.07, 15.26. C_22_H_23_N_2_ ESI-MS: m/z (M + H^+^): 315.2 (calculated), 315.2 (found).

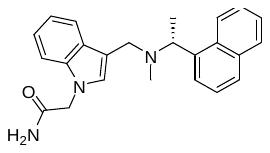

Jun9-85-2. Yield: 59%. ^1^H NMR (400 MHz, DMSO-d6) δ 8.42 (d, *J* = 8.4 Hz, 1H), 7.92 (d, *J* = 8.0 Hz, 1H), 7.83 (d, *J* = 8.0 Hz, 1H), 7.66 (d, *J* = 7.2 Hz, 1H), 7.57-7.42 (m, 4H), 7.36 (d, *J* = 8.0 Hz, 1H), 7.29 (d, *J* = 8.0 Hz, 1H), 7.22 (s, 1H), 7.20 (s, 1H), 7.09 (t, *J* = 7.6 Hz, 1H), 6.91 (t, *J* = 7.6 Hz, 1H), 4.74 (s, 2H), 4.47 (q, *J* = 6.4 Hz, 1H), 3.72 (d, *J* = 13.2 Hz, 1H), 3.66 (d, *J* = 13.2 Hz, 1H), 2.13 (s, 3H), 1.54 (d, *J* = 6.4 Hz, 3H). ^13^C NMR (100 MHz, DMSO-d6) δ 169.54, 140.43, 136.76, 133.66, 131.46, 129.03, 128.37, 127.83, 127.21, 125.41, 125.37, 125.22, 124.80, 124.36, 121.03, 119.27, 118.48, 111.61, 109.54, 58.78, 49.17, 48.29, 37.81, 15.17. C_24_H_26_N_3_O ESI-MS: m/z (M + H^+^): 372.2 (calculated), 372.2 (found).

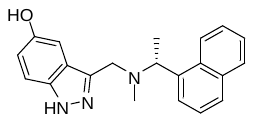

Jun9-86-1. Yield: 44%. ^1^H NMR (400 MHz, DMSO-d6) δ 12.50 (s, 1H), 8.98 (s, 1H), 8.34 (d, *J* = 8.4 Hz, 1H), 7.90 (d, *J* = 7.6 Hz, 1H), 7.82 (d, *J* = 8.0 Hz, 1H), 7.67 (d, *J* = 7.2 Hz, 1H), 7.55-7.40 (m, 3H), 7.31 (d, *J* = 8.8 Hz, 1H), 7.03 (d, *J* = 2.0 Hz, 1H), 6.90 (dd, *J* = 8.8, 2.0 Hz, 1H), 4.45 (q, *J* = 6.4 Hz, 1H), 3.91 (d, *J* = 13.2 Hz, 1H), 3.84 (d, *J* = 13.2 Hz, 1H), 2.07 (s, 3H), 1.55 (d, *J* = 6.4 Hz, 3H). ^13^C NMR (100 MHz, DMSO-d6) δ 150.83, 141.56, 140.20, 136.27, 133.59, 131.35, 128.40, 127.22, 125.52, 125.37, 125.28, 124.46, 124.39, 122.70, 117.57, 110.53, 102.68, 58.57, 51.17, 38.04, 15.86. C_21_H_22_N_3_O ESI-MS: m/z (M + H^+^): 332.2 (calculated), 332.2 (found).

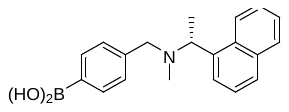

Jun9-84-4. Yield: 55%. ^1^H NMR (400 MHz, DMSO-d6) δ 8.49-8.39 (m, 1H), 7.92-7.47 (m, 8H), 7.25-7.13 (m, 2H), 4.55-4.37 (m, 1H), 3.63 (d, *J* = 8.0 Hz, 1H), 3.42 (d, *J* = 8.0 Hz, 1H), 2.09 (s, 3H), 1.51 (s, 3H). ^13^C NMR (100 MHz, DMSO-d6) δ 142.01, 140.09, 133.80, 133.68, 131.39, 128.48, 127.48, 127.36, 125.50, 125.45, 125.41, 125.25, 124.62, 124.33, 59.42, 58.07, 37.74, 15.40. C_20_H_23_BNO_2_ ESI-MS: m/z (M + H^+^): 320.2 (calculated), 320.2 (found).

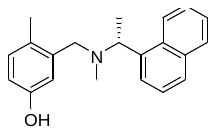

Jun9-87-1. Yield: 66%. ^1^H NMR (400 MHz, DMSO-d6) δ 9.02 (s, 1H), 8.26 (d, *J* = 8.0 Hz, 1H), 7.95-7.85 (m, 1H), 7.81 (d, *J* = 8.0 Hz, 1H), 7.62 (d, *J* = 7.2 Hz, 1H), 7.56-7.43 (m, 3H), 6.88 (d, *J* = 8.0 Hz, 1H), 6.77 (d, *J* = 2.4 Hz, 1H), 6.53 (dd, *J* = 8.0, 2.4 Hz, 1H), 4.46 (q, *J* = 6.4 Hz, 1H), 3.51 (d, *J* = 13.2 Hz, 1H), 3.37 (d, *J* = 13.2 Hz, 1H), 2.07 (s, 3H), 1.99 (s, 3H), 1.51 (d, *J* = 6.4 Hz, 3H). ^13^C NMR (100 MHz, DMSO-d6) δ 155.10, 139.87, 138.22, 133.61, 131.52, 130.63, 128.41, 127.28, 126.44, 125.46, 125.36, 125.16, 124.39, 124.26, 116.31, 113.37, 58.57, 55.80, 37.94, 17.81, 14.68. C_21_H_24_NO ESI-MS: m/z (M + H^+^): 306.2 (calculated), 306.2 (found).

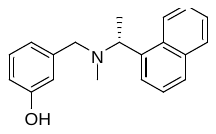

Jun9-75-2. Yield: 79%. ^1^H NMR (400 MHz, DMSO-d6) δ 8.45 (d, *J* = 8.4 Hz, 1H), 7.96-7.87 (m, 1H), 7.82 (d, *J* = 8.0 Hz, 1H), 7.62 (d, *J* = 7.2 Hz, 1H), 7.59-7.43 (m, 3H), 7.03 (t, *J* = 7.6 Hz, 1H), 6.68 (s, 1H), 6.64-6.56 (m, 2H), 4.42 (q, *J* = 6.4 Hz, 1H), 3.52 (d, *J* = 13.2 Hz, 1H), 3.33 (d, *J* = 13.2 Hz, 1H), 2.07 (s, 3H), 1.48 (d, *J* = 6.4 Hz, 3H). ^13^C NMR (100 MHz, DMSO-d6) δ 157.69, 141.19, 140.12, 133.67, 131.40, 128.89, 128.45, 127.29, 125.55, 125.42, 125.23, 124.56, 124.27, 118.61, 115.33, 113.75, 59.41, 58.33, 37.69, 15.56. C_20_H_22_NO ESI-MS: m/z (M + H^+^): 292.2 (calculated), 292.2 (found).

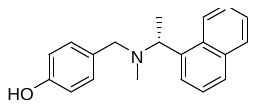

Jun9-72-2. Yield: 83%. ^1^H NMR (400 MHz, DMSO-d6) δ 9.22 (s, 1H), 8.42 (d, *J* = 8.0 Hz, 1H), 7.96-7.87 (m, 1H), 7.81 (d, *J* = 8.0 Hz, 1H), 7.61 (d, *J* = 7.2 Hz, 1H), 7.57-7.43 (m, 3H), 7.00 (d, *J* = 8.4 Hz, 2H), 6.66 (d, *J* = 8.4 Hz, 2H), 4.41 (q, *J* = 6.4 Hz, 1H), 3.49 (d, *J* = 12.8 Hz, 1H), 3.30 (d, *J* = 12.8 Hz, 1H), 2.04 (s, 3H), 1.48 (d, *J* = 6.4 Hz, 3H). ^13^C NMR (100 MHz, DMSO-d6) δ 156.12, 140.20, 133.69, 131.42, 129.71, 129.57, 128.47, 127.29, 125.48, 125.43, 125.26, 124.63, 124.32, 114.84, 59.15, 57.71, 37.38, 15.34. C_20_H_22_NO ESI-MS: m/z (M + H^+^): 292.2 (calculated), 292.2 (found).

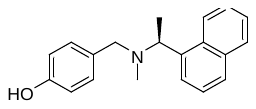

(*S*)-N-methyl-1-(naphthalen-1-yl)ethan-1-amine was used.

Jun9-84-5. Yield: 80%. ^1^H NMR (400 MHz, DMSO-d6) δ 9.21 (s, 1H), 8.42 (d, *J* = 8.0 Hz, 1H), 7.97-7.87 (m, 1H), 7.81 (d, *J* = 8.0 Hz, 1H), 7.61 (d, *J* = 7.2 Hz, 1H), 7.57-7.44 (m, 3H), 7.01 (d, *J* = 8.4 Hz, 2H), 6.66 (d, *J* = 8.4 Hz, 2H), 4.41 (q, *J* = 6.4 Hz, 1H), 3.49 (d, *J* = 12.8 Hz, 1H), 3.30 (d, *J* = 12.8 Hz, 1H), 2.04 (s, 3H), 1.48 (d, *J* = 6.4 Hz, 3H). ^13^C NMR (100 MHz, DMSO-d6) δ 156.11, 140.18, 133.67, 131.39, 129.67, 129.54, 128.44, 127.26, 125.44, 125.40, 125.22, 124.61, 124.29, 114.82, 59.14, 57.69, 37.35, 15.32. C_20_H_22_NO ESI-MS: m/z (M + H^+^): 292.2 (calculated), 292.2 (found).

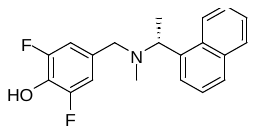

Jun9-86-2Yield: 80%. ^1^H NMR (400 MHz, DMSO-d6) δ 9.90 (s, 1H), 8.42 (d, *J* = 8.4 Hz, 1H), 8.02-7.89 (m, 1H), 7.82 (d, *J* = 8.0 Hz, 1H), 7.68-7.39 (m, 4H), 6.81 (d, *J* = 8.8 Hz, 2H), 4.48 (q, *J* = 6.4 Hz, 1H), 3.48 (d, *J* = 13.6 Hz, 1H), 3.32 (d, *J* = 13.6 Hz, 1H), 2.09 (s, 3H), 1.48 (d, *J* = 6.4 Hz, 3H). ^13^C NMR (100 MHz, DMSO-d6) δ 153.27 (d, *J* = 7.0 Hz), 150.87 (d, *J* = 7.0 Hz), 139.70, 133.69, 131.95, 131.38, 130.94, 128.51, 127.44, 125.50, 125.47, 125.21, 124.57, 124.28, 111.14 (dd, *J* = 15.0, 7.0 Hz), 59.04, 56.50, 37.56, 14.83. C_20_H_20_F_2_NO ESI-MS: m/z (M + H^+^): 328.2 (calculated), 328.2 (found).

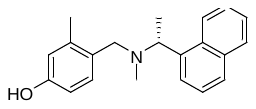

Jun9-87-2. Yield: 65%. ^1^H NMR (400 MHz, DMSO-d6) δ 8.19 (d, *J* = 8.0 Hz, 1H), 7.92-.84 (m, 1H), 7.80 (d, *J* = 8.0 Hz, 1H), 7.59 (d, *J* = 7.2 Hz, 1H), 7.50-7.41 (m, 3H), 7.01 (d, *J* = 8.0 Hz, 1H), 6.60-6.48 (m, 2H), 4.44 (q, *J* = 6.4 Hz, 1H), 3.51 (d, *J* = 12.8 Hz, 1H), 3.34 (d, *J* = 12.8 Hz, 1H), 2.02 (s, 6H), 1.50 (d, *J* = 6.4 Hz, 3H). ^13^C NMR (100 MHz, DMSO-d6) δ 156.15, 139.89, 138.16, 133.67, 131.62, 130.99, 128.80, 128.40, 127.28, 125.34, 125.14, 124.52, 124.34, 116.90, 112.10, 58.09, 55.22, 37.57, 18.84, 13.91. C_21_H_24_NO ESI-MS: m/z (M + H^+^): 306.2 (calculated), 306.2 (found).

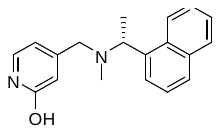

Jun9-84-6. Yield: 74%. ^1^H NMR (400 MHz, DMSO-d6) δ 8.42 (d, *J* = 8.4 Hz, 1H), 7.96-7.89 (m, 1H), 7.83 (d, *J* = 8.0 Hz, 1H), 7.62-7.45 (m, 4H), 7.20 (d, *J* = 6.8 Hz, 1H), 6.18 (s, 1H), 5.98 (dd, *J* = 6.8, 1.2 Hz, 1H), 4.48 (q, *J* = 6.4 Hz, 1H), 3.39 (d, *J* = 14.4 Hz, 1H), 3.25 (d, *J* = 14.4 Hz, 1H), 2.13 (s, 3H), 1.48 (d, *J* = 6.4 Hz, 3H). ^13^C NMR (100 MHz, DMSO-d6) δ 162.50, 154.05, 139.60, 134.56, 133.68, 131.39, 128.51, 127.50, 125.61, 125.49, 125.22, 124.50, 124.29, 117.64, 105.20, 59.36, 56.93, 38.01, 15.04. C_19_H_21_N_2_O ESI-MS: m/z (M + H^+^): 293.2 (calculated), 293.2 (found).

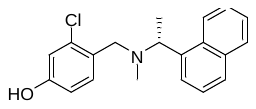

Jun9-87-3. Yield: 71%. ^1^H NMR (400 MHz, DMSO-d6) δ 8.37 (dd, *J* = 6.0, 3.6 Hz, 1H), 7.90 (dd, *J* = 6.0, 3.6 Hz, 1H), 7.81 (d, *J* = 8.0 Hz, 1H), 7.60 (d, *J* = 7.2 Hz, 1H), 7.55-7.41 (m, 3H), 7.19 (d, *J* = 8.4 Hz, 1H), 6.77 (d, *J* = 2.4 Hz, 1H), 6.68 (dd, *J* = 8.4, 2.4 Hz, 1H), 4.49 (q, *J* = 6.4 Hz, 1H), 3.56 (d, *J* = 13.6 Hz, 1H), 3.52 (d, *J* = 13.6 Hz, 1H), 2.06 (s, 3H), 1.50 (d, *J* = 6.4 Hz, 3H). ^13^C NMR (100 MHz, DMSO-d6) δ 156.97, 139.81, 133.64, 133.36, 131.58, 131.47, 128.41, 127.36, 126.71, 125.47, 125.41, 125.20, 124.61, 124.31, 115.58, 114.29, 59.01, 54.91, 37.50, 14.93. C_20_H_21_ClNO ESI-MS: m/z (M + H^+^): 326.2 (calculated), 326.2 (found).

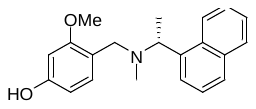

Jun9-75-5. Yield: 75%. ^1^H NMR (400 MHz, DMSO-d6) δ 9.26 (s, 1H), 8.48-8.35 (m, 1H), 7.95-7.85 (m, 1H), 7.79 (d, *J* = 8.0 Hz, 1H), 7.59 (d, *J* = 7.2 Hz, 1H), 7.55-7.41 (m, 3H), 7.00 (d, *J* = 8.0 Hz, 1H), 6.36 (d, *J* = 2.0 Hz, 1H), 6.29 (dd, *J* = 8.0, 2.0 Hz, 1H), 4.43 (q, *J* = 6.4 Hz, 1H), 3.66 (s, 3H), 3.50 (d, *J* = 13.2 Hz, 1H), 3.37 (d, *J* = 13.2 Hz, 1H), 2.04 (s, 3H), 1.48 (d, *J* = 6.4 Hz, 3H). ^13^C NMR (100 MHz, DMSO-d6) δ 158.30, 157.42, 140.23, 133.61, 131.54, 130.46, 128.30, 127.16, 125.33, 125.30, 125.15, 124.81, 124.20, 117.27, 106.43, 98.63, 59.05, 54.89, 51.99, 37.14, 14.77. C_21_H_24_NO_2_ ESI-MS: m/z (M + H^+^): 322.2 (calculated), 322.2 (found).

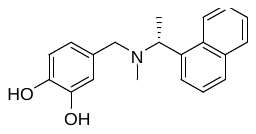

Jun9-75-3. Yield: 70%. ^1^H NMR (400 MHz, DMSO-d6) δ 8.49-8.22 (m, 1H), 8.09-7.28 (m, 9H), 6.38-6.25 (m, 2H), 4.56 (q, *J* = 6.4 Hz, 1H), 3.44 (d, *J* = 12.4 Hz, 1H), 3.24 (dd, *J* = 12.4, 3.2 Hz, 1H), 2.29 (s, 3H), 1.41 (d, *J* = 6.4 Hz, 3H). ^13^C NMR (100 MHz, DMSO-d6) δ 153.11, 151.92, 140.66, 134.00, 131.39, 129.18, 127.52, 126.37, 126.11, 125.89, 125.67, 123.53, 123.31, 117.84, 108.72, 106.93, 59.28, 55.33, 34.28, 23.30. C_20_H_22_NO_2_ ESI-MS: m/z (M + H^+^): 308.2 (calculated), 308.2 (found).

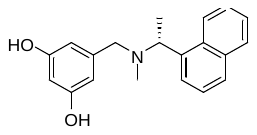

Jun9-67-1. Yield: 73%. ^1^H NMR (400 MHz, DMSO-d6) δ 9.06 (s, 2H), 8.45 (d, *J* = 8.0 Hz, 1H), 7.92 (d, *J* = 8.0 Hz, 1H), 7.82 (d, *J* = 8.0 Hz, 1H), 7.62 (d, *J* = 7.2 Hz, 1H), 7.58-7.45 (m, 3H), 6.15 (d, *J* = 2.0 Hz, 2H), 6.06 (t, *J* = 2.0 Hz, 1H), 4.40 (q, *J* = 6.4 Hz, 1H), 3.44 (d, *J* = 13.2 Hz, 1H), 3.24 (d, *J* = 13.2 Hz, 1H), 2.07 (s, 3H), 1.47 (d, *J* = 6.4 Hz, 3H). ^13^C NMR (100 MHz, DMSO-d6) δ 158.14, 141.89, 140.20, 133.67, 131.40, 128.47, 127.29, 125.62, 125.44, 125.25, 124.54, 124.26, 106.37, 100.97, 59.47, 58.62, 37.77, 15.78. C_20_H_22_NO_2_ ESI-MS: m/z (M + H^+^): 308.2 (calculated), 308.2 (found).

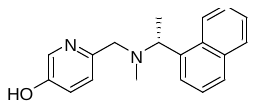

Jun9-86-8. Yield: 72%. ^1^H NMR (400 MHz, DMSO-d6) δ 8.41 (d, *J* = 8.0 Hz, 1H), 8.02 (d, *J* = 2.4 Hz, 1H), 7.96-7.86 (m, 1H), 7.81 (d, *J* = 8.0 Hz, 1H), 7.61 (d, *J* = 7.2 Hz, 1H), 7.59-7.44 (m, 3H), 7.16-7.05 (m, 2H), 4.47 (q, *J* = 6.4 Hz, 1H), 3.64 (d, *J* = 13.6 Hz, 1H), 3.51 (d, *J* = 13.6 Hz, 1H), 2.10 (s, 3H), 1.48 (d, *J* = 6.4 Hz, 3H). ^13^C NMR (100 MHz, DMSO-d6) δ 152.30, 149.99, 140.01, 136.43, 133.68, 131.44, 128.47, 127.33, 125.54, 125.41, 125.24, 124.55, 124.31, 122.88, 122.70, 59.38, 59.11, 37.96, 15.39. C_19_H_21_N_2_O ESI-MS: m/z (M + H^+^): 293.2 (calculated), 293.2 (found).

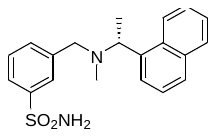

Jun9-86-5. Yield: 59%. ^1^H NMR (400 MHz, DMSO-d6) δ 8.45 (d, *J* = 8.4 Hz, 1H), 7.93 (d, *J* = 7.8 Hz, 1H), 7.83 (d, *J* = 8.0 Hz, 1H), 7.78 (s, 1H), 7.70-7.64 (m, 2H), 7.60-7.40 (m, 5H), 7.32 (s, 2H), 4.52 (q, *J* = 6.4 Hz, 1H), 3.71 (d, *J* = 13.6 Hz, 1H), 3.53 (d, *J* = 13.6 Hz, 1H), 2.09 (s, 3H), 1.52 (d, *J* = 6.4 Hz, 3H). ^13^C NMR (100 MHz, DMSO-d6) δ 144.14, 140.98, 139.76, 133.68, 131.60, 131.39, 128.78, 128.51, 127.45, 125.72, 125.51, 125.30, 125.26, 124.46, 124.35, 124.16, 59.31, 57.85, 37.66, 15.63. C_20_H_23_N_2_O_2_S ESI-MS: m/z (M + H^+^): 355.2 (calculated), 355.2 (found).

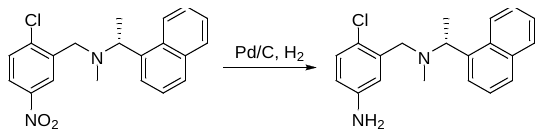

The crude product of the reductive amination was dissolved in ethyl acetate (10 mL) and 10% Pd/C (50 mg) was added. The reaction was stirred under H_2_ balloon until TLC showed the starting material was completely consumed. The reaction was filtered and washed with CH_2_Cl_2_/MeOH (CH_2_Cl_2_/MeOH = 15:1). The combined organic layer was concentrated under reduced pressure. The residue was purified by silica gel flash column chromatography (CH_2_Cl_2_ to CH_2_Cl_2_/MeOH = 10:1) to afford the target product.

Jun9-81-3. Yield: 71%. ^1^H NMR (400 MHz, DMSO-d6) δ 8.40 (d, *J* = 8.0 Hz, 1H), 7.97-7.85 (m, 1H), 7.81 (d, *J* = 8.0 Hz, 1H), 7.64 (d, *J* = 7.2 Hz, 1H), 7.57-7.42 (m, 3H), 6.98 (d, *J* = 8.4 Hz, 1H), 6.77 (d, *J* = 2.8 Hz, 1H), 6.44 (dd, *J* = 8.4, 2.8 Hz, 1H), 5.17 (s, 2H), 4.47 (q, *J* = 6.4 Hz, 1H), 3.53 (d, *J* = 14.0 Hz, 1H), 3.48 (d, *J* = 14.0 Hz, 1H), 2.11 (s, 3H), 1.50 (d, *J* = 6.4 Hz, 3H). ^13^C NMR (100 MHz, DMSO-d6) δ 147.68, 140.00, 136.64, 133.60, 131.36, 129.17, 128.42, 127.29, 125.65, 125.42, 125.25, 124.42, 124.23, 118.96, 115.55, 113.80, 59.28, 55.55, 38.02, 16.10. C_20_H_22_ClN_2_ ESI-MS: m/z (M + H^+^): 325.2 (calculated), 325.2 (found).

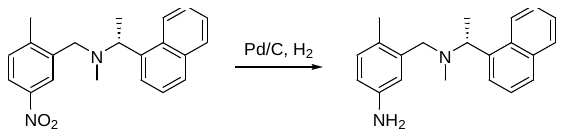

Following the procedure for Jun9-81-3.

Jun9-53-2. Yield: 30%. ^1^H NMR (400 MHz, DMSO-d6) δ 8.28-8.19 (m, 1H), 7.93-7.86 (m, 1H), 7.81 (d, *J* = 8.0 Hz, 1H), 7.62 (d, *J* = 7.2 Hz, 1H), 7.51-7.43 (m, 3H), 6.77 (d, *J* = 8.0 Hz, 1H), 6.60 (d, *J* = 2.4 Hz, 1H), 6.38 (dd, *J* = 8.0, 2.4 Hz, 1H), 4.75 (s, 2H), 4.44 (q, *J* = 6.4 Hz, 1H), 3.49 (d, *J* = 12.8 Hz, 1H), 3.32 (d, *J* = 12.88 Hz, 1H), 2.06 (s, 3H), 1.96 (s, 3H), 1.51 (d, *J* = 6.4 Hz, 3H). ^13^C NMR (100 MHz, DMSO-d6) δ 146.13, 139.95, 137.30, 133.59, 131.52, 130.32, 128.36, 127.22, 125.41, 125.33, 125.15, 124.43, 124.27, 123.59, 115.80, 112.61, 58.33, 56.27, 37.92, 17.83, 14.66. C_21_H_25_N_2_ ESI-MS: m/z (M + H^+^): 305.2 (calculated), 305.2 (found).

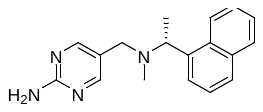

Jun9-85-5. Yield: 27%. ^1^H NMR (400 MHz, DMSO-d6) δ 8.40 (d, *J* = 8.0 Hz, 1H), 7.97 (s, 2H), 7.96-7.89 (m, 1H), 7.83 (d, *J* = 8.0 Hz, 1H), 7.60 (d, *J* = 7.2 Hz, 1H), 7.57-7.46 (m, 3H), 6.47 (s, 2H), 4.44 (q, *J* = 6.4 Hz, 1H), 3.40 (d, *J* = 13.2 Hz, 1H), 3.24 (d, *J* = 13.2 Hz, 1H), 2.07 (s, 3H), 1.48 (d, *J* = 6.4 Hz, 3H). ^13^C NMR (100 MHz, DMSO-d6) δ 162.92, 158.19, 139.82, 133.69, 131.32, 128.50, 127.40, 125.49, 125.44, 125.24, 124.55, 124.37, 120.13, 58.98, 52.44, 37.03, 15.05. C_18_H_21_N_4_ ESI-MS: m/z (M + H^+^): 293.2 (calculated), 293.2 (found).

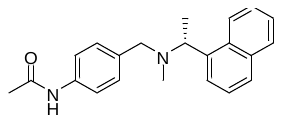

Jun9-85-6. Yield: 50%. ^1^H NMR (400 MHz, DMSO-d6) δ 9.86 (s, 1H), 8.43 (d, *J* = 8.4 Hz, 1H), 7.92 (d, *J* = 7.6 Hz, 1H), 7.82 (d, *J* = 8.0 Hz, 1H), 7.62 (d, *J* = 7.2 Hz, 1H), 7.59-7.44 (m, 5H), 7.12 (d, *J* = 8.4 Hz, 2H), 4.43 (q, *J* = 6.4 Hz, 1H), 3.54 (d, *J* = 13.2 Hz, 1H), 3.36 (d, *J* = 13.2 Hz, 1H), 2.06 (s, 3H), 2.03 (s, 3H), 1.49 (d, *J* = 6.4 Hz, 3H). ^13^C NMR (100 MHz, DMSO-d6) δ 168.05, 140.06, 137.95, 134.21, 133.67, 131.38, 128.64, 128.46, 127.31, 125.49, 125.42, 125.23, 124.59, 124.30, 118.74, 59.24, 57.68, 37.50, 23.92, 15.36. C_22_H_25_N_2_O ESI-MS: m/z (M + H^+^): 333.2 (calculated), 333.2 (found).

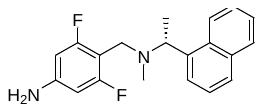

Jun9-81-2. Yield: 49%. ^1^H NMR (400 MHz, DMSO-d6) δ 8.29 (d, *J* = 8.4 Hz, 1H), 7.93-7.84 (m, 1H), 7.80 (d, *J* = 8.0 Hz, 1H), 7.57 (d, *J* = 6.8 Hz, 1H), 7.49-7.35 (m, 2H), 6.30-6.15 (m, 3H), 5.66 (s, 2H), 4.41 (q, *J* = 6.4 Hz, 1H), 3.53 (d, *J* = 12.4 Hz, 1H), 3.45 (d, *J* = 12.4 Hz, 1H), 1.97 (s, 3H), 1.46 (d, *J* = 6.4 Hz, 3H). ^13^C NMR (100 MHz, DMSO-d6) δ 163.29, 161.47, 150.27, 139.78, 133.58, 131.48, 128.31, 127.23, 125.32, 125.28, 125.13, 124.47, 124.22, 100.40, 100.24, 96.08, 95.86, 58.44, 45.21, 36.38, 14.69. C_20_H_21_F_2_N_2_ ESI-MS: m/z (M + H^+^): 327.2 (calculated), 327.2 (found).

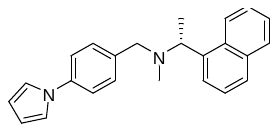

Jun9-67-5. Yield: 48%. ^1^H NMR (400 MHz, DMSO-d6) δ 8.46 (d, *J* = 8.4 Hz, 1H), 7.93 (d, *J* = 8.0 Hz, 1H), 7.83 (d, *J* = 8.0 Hz, 1H), 7.63 (d, *J* = 7.2 Hz, 1H), 7.61-7.47 (m, 2H), 7.45 (d, *J* = 8.4 Hz, 2H), 7.33-7.28 (m, 2H), 7.26 (d, *J* = 8.4 Hz, 2H), 6.32-6.16 (m, 2H), 4.48 (q, *J* = 6.4 Hz, 1H), 3.60 (d, *J* = 13.2 Hz, 1H), 3.44 (d, *J* = 13.2 Hz, 1H), 2.10 (s, 3H), 1.51 (d, *J* = 6.4 Hz, 3H). ^13^C NMR (100 MHz, DMSO-d6) δ 139.95, 138.63, 136.84, 133.71, 131.43, 129.49, 128.51, 127.41 , 125.55, 125.46, 125.24, 124.61, 124.33, 119.06, 118.86, 110.24, 59.32, 57.29, 37.60, 15.15. C_24_H_25_N_2_ ESI-MS: m/z (M + H^+^): 341.2 (calculated), 341.2 (found).

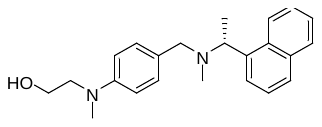

Jun9-86-4. Jun9-67-5. Yield: 57%. ^1^H NMR (400 MHz, DMSO-d6) δ 8.43 (d, *J* = 8.0 Hz, 1H), 7.96-7.87 (m, 1H), 7.81 (d, *J* = 8.0 Hz, 1H), 7.61 (d, *J* = 7.2 Hz, 1H), 7.58-7.45 (m, 3H), 7.00 (d, *J* = 8.4 Hz, 2H), 6.59 (d, *J* = 8.4 Hz, 2H), 4.40 (q, *J* = 6.4 Hz, 1H), 3.57-3.44 (m, 3H), 3.39-3.25 (m, 3H), 2.88 (s, 3H), 2.05 (s, 3H), 1.48 (d, *J* = 6.4 Hz, 3H). ^13^C NMR (100 MHz, DMSO-d6) δ 148.12, 140.27, 133.68, 131.42, 129.34, 128.43, 127.24, 126.12, 125.40, 125.39, 125.21, 124.72, 124.30, 111.39, 59.11, 58.04, 54.89, 54.39, 38.62, 37.29, 15.19. C_23_H_29_N_2_O ESI-MS: m/z (M + H^+^): 349.2 (calculated), 349.2 (found).
